# Computational screen of promoter configurations that robustly sense transcription factor dynamics

**DOI:** 10.64898/2026.05.19.726340

**Authors:** T. Curtis Shoyer, Barbara Di Ventura

## Abstract

Transcription factors (TFs) respond to external stimuli with time-varying changes in activity or localization (TF dynamics), driving differential transcriptional programs. Previous studies indicated that TF dynamics can be decoded at the promoter level in eukaryotes, yet a systematic understanding of robust solutions is lacking. By computationally screening over 10,000 mathematical models of multi-state promoters with various forms of TF-mediated regulation, we identify robust configurations that selectively respond to sustained (“pulse filtering”) or pulsatile (“pulse boosting”) TF dynamics. Promoters that activate via intermediate states and have negatively regulated deactivation robustly perform pulse filtering. In contrast, robust pulse boosting is achieved by promoters with a TF-mediated refractory state that permits short activation and recovers between pulses. Bifunctional TFs that exert activator- and repressor-like regulation extend the design space for pulse boosting. These results reveal general principles by which promoters interpret TF dynamics and suggest strategies to engineer synthetic systems to exploit them.

**Highlights:**

- Computational screen of over 10,000 promoter models identifies features that enable promoters to selectively respond to sustained (“pulse filtering”) or pulsatile (“pulse boosting”) transcription factor (TF) dynamics.
- Promoters that activate via intermediate states and have negatively regulated deactivation robustly perform pulse filtering.
- Promoters with TF-regulated refractoriness robustly perform pulse boosting.
- Promoters regulated by bifunctional TFs extend the design space for pulse boosting.

## Introduction

Cells use transcription factors (TFs) to transduce stimuli into gene expression outputs. To do so, TFs bind to regulatory regions upstream of their target genes and modulate their transcriptional activity. In many signaling pathways, TF activity varies over time through regulated changes in TF abundance, localization, or post-translational modification (1,2). These TF dynamics have typical timescales of one minute to several hours (1,2) and should be distinguished from the often more rapid DNA binding dynamics (3). TF dynamics can encode information about the identity and strength of stimuli and are linked to different gene expression programs and cellular outcomes (1,2). For example, levels of the tumor suppressor p53 exhibit multiple pulses in response to γ-radiation versus a single sustained pulse in response to UV-radiation, and these different p53 dynamics result in different cell fates (4). Stimulus-specific dynamics have also been observed for NF-κB in inflammation and immune response (5,6), Erk in proliferation and differentiation (7–9), and the yeast Msn2 and Crz1 in stress responses (10–12). While the upstream mechanisms that generate TF dynamics are well established, how cells decode TF dynamics to control gene expression remains more challenging.

Mathematical modeling has played a central role in understanding how cells decode dynamic signals. In gene regulatory networks, feedforward and feedback motifs comprised of multiple interacting genes or proteins have been shown to process temporal signals (13–18). Computational screens have identified core network features that robustly perform these functions over a wide range of parameters consistent with biological variations (13,14,16). For instance, transcriptional networks composed of a TF that regulates a target gene both directly and via an intermediate gene can selectively respond to sustained or pulsatile TF inputs, depending on the logic and regulation of the branches (14,17). These network motifs often make use of branches driven by different biochemical processes and occurring at different timescales and/or cellular compartments (1,16). For example, a coherent feedforward loop in which ERK both activates transcription of c-Fos and phosphorylates accumulated c-Fos turns on pro-differentiation genes in response to sustained but not transient ERK signals (9,16).

While strategies for temporal processing by gene networks involve the co-regulation of a target gene by multiple interacting genes, it is also possible that a single promoter is directly responsive to TF dynamics. Indeed, mathematical models have been proposed for the decoding of TF dynamics at the level of the target promoter itself (11,12,19–27). These models have been developed to understand how the properties of target promoters affect their sensitivity to TF dynamics and have been experimentally tested by directly manipulating the dynamics of natural or synthetic TFs via chemical or optogenetic approaches (11,12,23–25,27–29) (Table S1). The core concept of these models is that promoters transition between a set of distinct biochemical states, and that the dynamics of these transitions are regulated by bound TFs and co-effectors to control the transcriptional output. Notably, the lifetimes of key promoter states and associated mechanisms exhibit a range of timescales (roughly 10 seconds to 100 minutes) (12,27,28,30,31) that overlap those typical of TF dynamics (1 minute to several hours) (1,2). Promoter models differ in the number of states, the allowed state transitions, and the way in which TFs regulate transition kinetics. The simplest models contain two states: inactive (non-transcribing) and active (transcribing). More complex models incorporate additional non-transcribing states, such as primed states (12,24–26) that are accessed prior to activation and often associated with chromatin remodeling (12,26,32), or refractory states accessed after activation, in which the promoter is temporarily non-responsive even if the TF is bound (23,28,30,33).

Different promoter models have been used to explain the responses of different target genes. Models with one or several primed states, tied to chromatin-associated processes with relatively long timescales, can explain how some Msn2, Crz1, and NF-κB target genes respond selectively to sustained, longer duration TF inputs compared to one or multiple pulses of shorter durations (12,24,26,32). We refer to this behavior as “pulse filtering” (which resembles a low-pass filter in some capacity). In contrast, promoters that respond more to pulsatile than sustained TF dynamics, which we refer to as “pulse boosting”, have been observed less often. One example is the Crz1-responsive pYPS1 promoter, which could be explained by a two-state model that required an easily saturable dose response, fast activation, and slow inactivation (24).

Taken together, different models have been applied to specific TFs and target genes, often under differing experimental conditions, such as different input dynamics and readouts that cannot be directly compared. As such, the general design principles that enable promoters to discriminate between sustained and pulsatile inputs have remained so far unclear. Additionally, we lack a complete understanding of the TF-promoter design space, especially the list of possible solutions for pulse filtering and pulse boosting and a catalogue of TF and promoter features that enable robust decoding in the presence of biological variability and/or parameter uncertainty. A systematic understanding of this design space is needed both to explain how natural promoters decode signaling dynamics and to engineer synthetic systems with predictable temporal responses.

To address this gap, we use mathematical modeling to systematically investigate properties of promoter configurations (defined by the states, transitions, and types of regulation) that enable robust decoding of TF dynamics. By screening *in-silico* over 10,000 promoter configurations incorporating a variety of promoter states, transitions, and forms of TF-mediated regulation, we identify properties of promoter configurations and kinetic requirements that enable promoters to selectively respond to sustained or pulsatile TF dynamics. Configurations that activate via intermediate states and in which deactivation is negatively regulated by the TF robustly perform pulse filtering. These promoters operate in regimes where they activate slowly enough to prevent saturation during TF pulses and deactivate quickly enough to prevent accumulated activation between TF pulses. In contrast, configurations with a TF-mediated refractory state robustly perform pulse boosting. These promoters operate in regimes where a transient phase of high transcription is followed by refractoriness at longer timescales. Bifunctional TFs that exert activator- and repressor-like regulation extend the design space supporting pulse boosting, but in general promoter configurations are better suited to robustly perform pulse filtering than pulse boosting. Our results identify general design principles that enable promoters to robustly convert TF dynamics into transcriptional outputs, advancing a predictive understanding of how temporal signals are decoded into gene-expression programs and providing a roadmap for engineering synthetic TFs and promoters with defined temporal sensitivities.

## Results

### Modeling framework based on promoter states and TF-regulated promoter transitions

We constructed a mathematical model of gene expression based on a promoter that transitions between a set of distinct states and is subject to regulation by the TF (Figure 1A, Methods). We considered a variety of promoter states, transitions, and forms of TF regulation from existing promoter models (11,12,19–25,28). In our model, the sequence-specific TF binds to the enhancer (response elements) present upstream of the core promoter. We assume that TF:DNA binding/unbinding is sufficiently fast compared to other processes (including promoter state transitions), such that the bound fraction is at equilibrium (28). The equilibrium bound fraction is a function of the nuclear concentration of active TF, [TF] (Methods Eq. 1). The promoter transitions between two, three, or four possible states (Figure 1A). In the inactive state (I), mRNA cannot be transcribed, regardless if the TF is bound or not. In the active state (A), mRNA can be transcribed. This constitutes the simplest two-state model. Biologically, the transition to the A state involves the assembly of the pre-initiation complex (PIC) and associated processes (12). As many eukaryotic promoters sensitive to TF dynamics are best described with an intermediate state before activation (12,24–26,28), we also considered models with a primed state (P), which is transcriptionally inactive but closer to activation. If the P state is present, the promoter must transition from the I state through the P state to reach the A state. Biologically, entry into the P state can involve histone modifications and chromatin remodeling (12,32), as well as DNA looping (28). Furthermore, transcription from eukaryotic promoters occurs in bursts separated by intervals in which the promoter becomes temporarily non-responsive or refractory, motivating the inclusion of a non-transcriptional refractory state (R) (28,30,33–35). The R state is only reached from the A state. Biologically, refractoriness varies between genes (30) and can depend on aspects including core promoter, cis-regulatory elements, and chromatin remodeling (33–35). Importantly, the P and R states are distinct as they differ in the allowed transitions by which the promoter enters and exits them (Figure 1A). For simplicity, we considered at most one P and one R state, as has been done previously (12,23–25,28).

**Figure 1:**
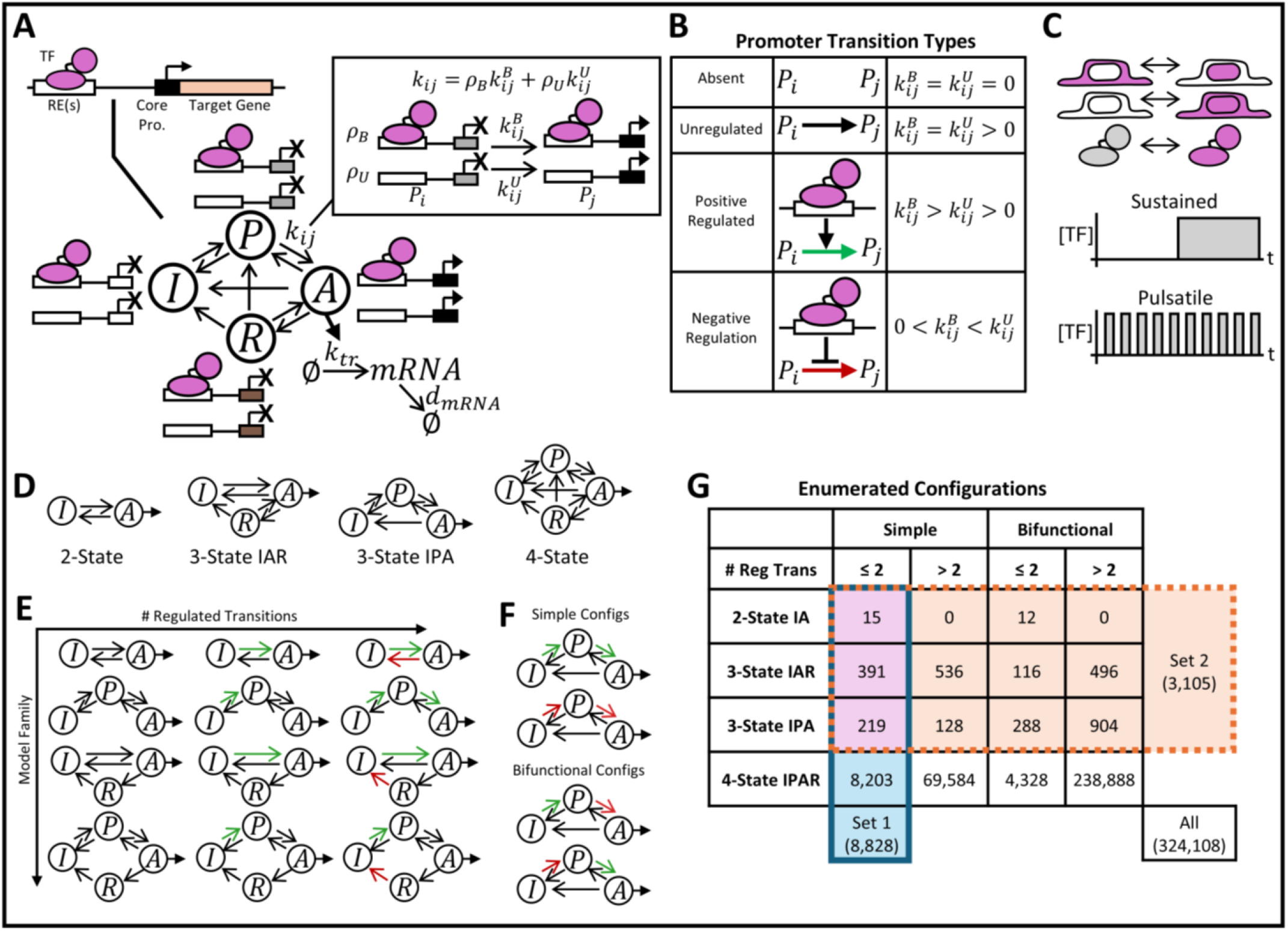
Model formulation and configuration enumeration. A) Model of a target promoter that transitions between states in a TF-dependent or -independent manner to regulate transcriptional output. All allowable state transitions for the 4-state model are depicted in black arrows. *k*_*ij*_ is the rate constant for the transition of the promoter from state *P*_*i*_ to *P*_*j*_, given in terms of rate constants for bound 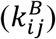 and unbound 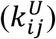 promoter, and the equilibrium bound (*ρ*_*B*_) and unbound (*ρ*_*U*_) fractions (Methods Eq. 1-3). *k*_*tr*_ is the transcription rate (Methods Eq. 4); *d*_*mRNA*_ is the mRNA degradation rate (Methods Eq. 5). B) Types of promoter state transitions defined by their 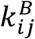 and 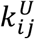 values (see Methods Eq. 3 for the definition of *k*_*ij*_ for each transition type). C) Model Input: Dynamics in the nuclear concentration of active TF, achieved by changes in TF localization, abundance, or activity. Sustained versus pulsatile TF dynamics with matched AUCs and amplitudes (Methods Eq. 7-8). D) Model families defined by their promoter states, each indicating all allowable state transitions in black arrows. E) Example configurations with varying numbers of TF-regulated transitions. TF-independent, positively regulated, or negatively regulated transitions are indicated in black, green, or red arrows, respectively. F) Examples of simple versus bifunctional configurations. G) Number of enumerated configurations in each classification category. Configuration sets 1 and 2 are indicated in blue and orange, respectively, with overlap in purple.

In our model, the TF can bind to the promoter in any state and regulate how the promoter transitions between states. We used a formalism for TF-regulated promoter state transitions from Li et al. (28) in which a transition can be positively regulated/accelerated by a bound TF, negatively regulated/decelerated by a bound TF, or be independent of the TF (Figure 1B, Methods Eq. 3). For example, in a 2-state model, a positively regulated I→A transition means the TF increases the activation rate constant when bound, and a negatively regulated A→I transition means the TF decreases the deactivation rate constant when bound. Transcription of mRNA occurs from the A state. We considered two transcription rate models commonly used in the literature (Methods Eq. 4): TF-independent, where transcription from the A state occurs at a constant rate regardless of bound TF (24,25,28), or TF-dependent, where transcription from the A state depends on the fraction of bound TF (12,23,28), being either TF-activated or TF-repressed. Together, this modeling framework can be used to formulate exact or approximate versions of a wide range of models proposed for promoters sensitive to TF dynamics and/or characterized by refractoriness (Table S1).

The input to the model is a time-dependent nuclear concentration of active TF (“TF dynamics”): [*TF*] = *f*(*t*). Based on the dynamics of natural TFs and previous experimental and theoretical work on the sensitivity of genes to TF dynamics (1,2,4,11,12,23–25), we compared the response to two TF dynamics: sustained (a single long square-wave pulse) and pulsatile (multiple shorter square-wave pulses) (Figure 1C; Methods Eq. 7-8). Importantly, we matched the area under the curve (AUC) and the amplitude (A) for [*TF*] across both conditions to exclude effects due to cumulative TF levels (4,23). Lastly, we describe the population-level dynamics of our model using a system of ordinary differential equations (ODEs) for the fractions of promoters in each state as well as the abundance of mRNA (Methods Eq. 5-6), as has been done previously (11,12,23–26,28).

### Enumeration and classification of promoter configurations

Taking inspiration from biological network enumeration and screening approaches (13,14,16,36,37), we systematically enumerated promoter configurations and screened them for the ability to robustly discriminate between sustained and pulsatile TF dynamics. We first enumerated all 2-, 3-, and 4-state promoter configurations that satisfied rules about state connectivity and biologically plausible state transitions, yielding over 324,000 configurations (Figure S1A-B). We further classified promoter configurations according to three properties: the available states (2-state IA, 3-State IAR, 3-State IPA, or 4-State IPAR; Figure 1D), the number of TF-regulated transitions (Figure 1E), and the type of TF-regulated transitions (Figure 1F). The latter distinguishes simple regulation, where the TF does *not* exert both activator-like and repressor-like regulation, from bifunctional regulation, where the TF does exert both (Figure S1C-E). We defined the following as activator-like: positive regulation of transitions on the path from I to A (I→A, I→P, P→A), negative regulation of the corresponding reverse transitions (I←A, I←P, P←A), and TF-dependent increase in transcription rate (Figure S1C). Conversely, we defined the following as repressor-like: negative regulation of forward transitions along the path from I to A (I→A, I→P, P→A), positive regulation of the corresponding reverse transitions (I←A, I←P, P←A), and TF-dependent decrease in transcription rate (Figure S1C). As there is incomplete evidence of how TFs influence transitions to/from the R state, all transitions to/from the R state were left unclassified, i.e. neither activator-nor repressor-like (Figure S1C).

As it was not feasible to screen all the over 324,000 enumerated configurations with sufficient parameter space coverage, we focused on two subsets. Configuration set 1 contains all 2-, 3-, and 4-state configurations with a maximum of 2 TF-regulated promoter transitions and in which the TF exerts simple regulation (Figure 1G and Figure S1F; 8,828 configurations in total). In support of this choice, this set includes exact or approximate versions of many models proposed for a range of promoters sensitive to TF dynamics and/or characterized by refractoriness (Table S1). To address the possibilities of bifunctional regulation and higher numbers of TF-regulated promoter transitions, configuration set 2 consists of all 2- and 3-state configurations, with no restrictions regarding the number or type of regulation exerted by the TF (Figure 1G and Figure S1G; 3,105 configurations in total, of which 625 overlap with configuration set 1).

### Screen of promoter configurations for the ability to discriminate TF dynamics

We systematically assessed each configuration for its ability to perform two specific functions over a large parameter space: pulse filtering and pulse boosting (Figure 2A). The lifetimes of key promoter states and associated mechanisms are estimated to span a large range of timescales (roughly 10 seconds to 100 minutes) (12,27,28,30,31) with respect to those typical for pulsatile TF dynamics (1 minute to several hours) (1,2). Thus, the parameter space varies the unbound rate constants of promoter transitions (*k*_*ij*,0_) across large ranges with respect to the timescales of pulsatile TF dynamics (*τ*_*on*_ and *τ*_*off*_). Variations in the strengths of TF regulation 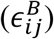, rate constants for mRNA transcription (*k*_*tr,max*_) and degradation (*d*_*mRNA*_), and TF binding parameters (*K*_*d,TF*_ and *n*_*TF*_) were also considered (see Table S2 for all parameter ranges and distributions). For each configuration we analyzed 1,000 parameter sets generated by Latin Hypercube Sampling (LHS). For each parameter set, we computed the steady-state mRNA in response to continuous TF absence ([*TF*] = 0) and presence ([*TF*] = *A*). Based on this comparison, we classified each parameter set as a steady-state activator (at least two-fold higher mRNA levels with TF present), repressor (at least two-fold lower mRNA levels with TF present), or non-responsive (Figure 2A). We considered parameter sets classified as activators for subsequent analyses. To assess sensitivity to TF dynamics, we analyzed the response to pulsatile and sustained TF inputs and computed a pulsatile-to-sustained ratio (PSR) using the mRNA AUC for each (Figure 2B). Parameter sets with more than two-fold higher gene expression for sustained (PSR < 1/2) or pulsatile (PSR > 2) inputs were classified as pulse filtering or pulse boosting, respectively (Figure 2A-B). In computational screens to evaluate the robustness of biological systems to parameter variations (e.g. due to heterogeneity, fluctuations, or uncertainty), robustness is often defined as the fraction of parameter sets for which a target function is performed above a threshold score (13,14,36). Therefore, for each promoter configuration we computed the pulse filtering/pulse boosting robustness as the fraction of parameter sets that perform activation and pulse filtering/pulse boosting.

**Figure 2:**
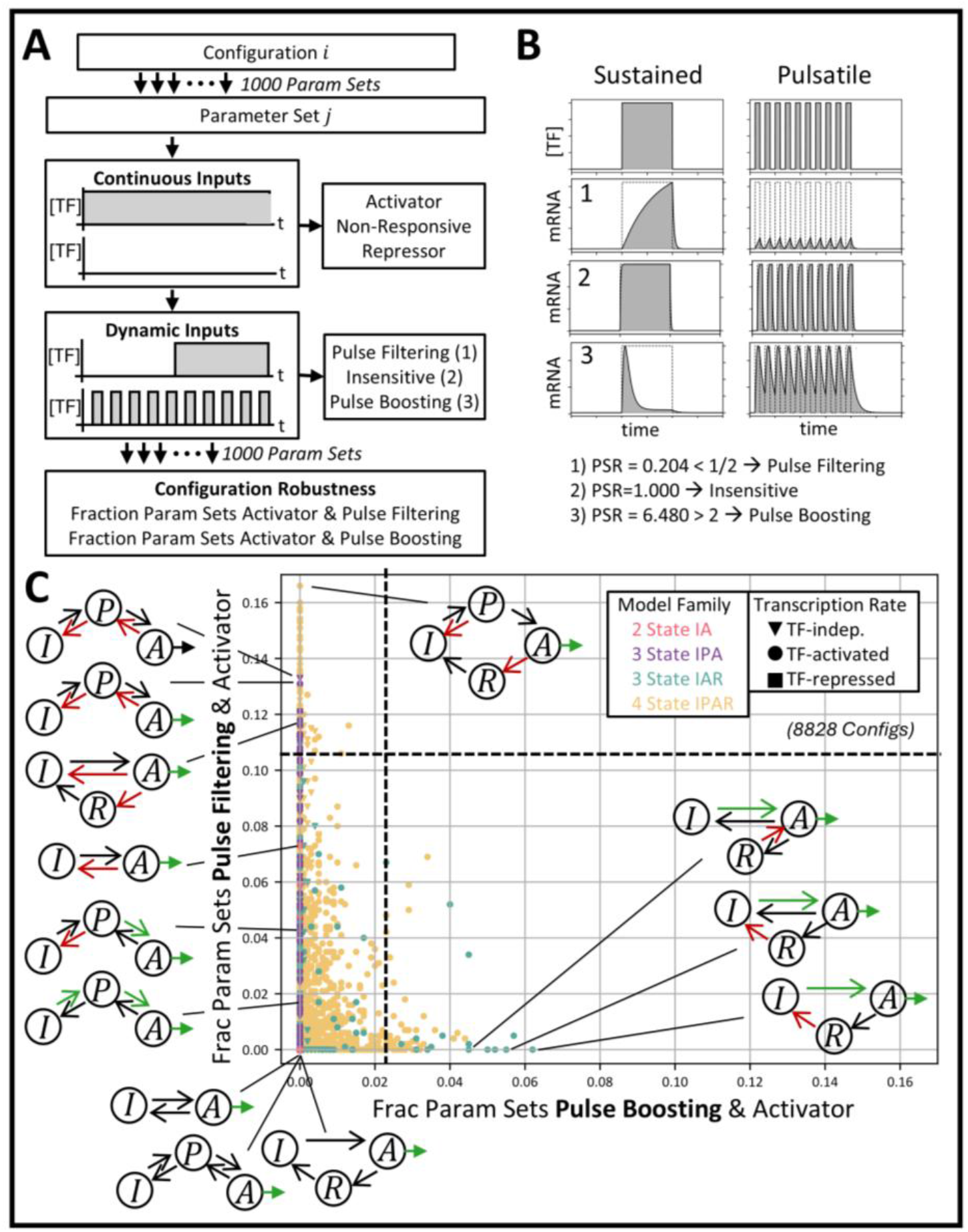
Screen of promoter configurations for the ability to discriminate TF dynamics. A) Overview of configuration screening workflow. B) Example time courses for sustained (left) and pulsatile (right) TF dynamics showing [TF] input followed by mRNA outputs for three different behaviors: (1) pulse filtering, (2) insensitivity to dynamics, and (3) pulse boosting. C) Response of each configuration in configuration set 1 to TF dynamics. Scatter plot of the fraction of parameter sets that are activators and perform pulse filtering (PSR<1/2) versus pulse boosting (PSR>2). Dotted lines correspond to cutoffs containing the top 1% most robust pulse filtering and pulse boosting configurations. See Figure S5 for data plotted separately by model family.

We first evaluated the configuration screening approach. We confirmed that screen results for both continuous and dynamic inputs with the selected parameter sample size (1,000 parameter sets) match those with a larger parameter sample size of 10,000 (Figure S2). We also verified that the results showed minimal variability across 10 independent replicates of the same sample size (Figure S2). Moreover, we tested screen results using a non-saturating versus saturating TF concentration (Figure S3). For both continuous and dynamic inputs, the robustness of configurations under saturating and non-saturating TF concentrations were strongly correlated. We therefore performed the screens under saturating TF conditions, where the TF-bound promoter fraction was approximately 1 in the presence of TF and 0 in the absence of TF. Lastly, we confirmed that screen results were robust to the PSR cutoff chosen for classifying parameter sets as pulse filtering and pulse boosting (Figure S4).

We applied the screening approach to configuration set 1, comprised of all 2-, 3-, and 4-state configurations with simple regulation and at most 2 TF-regulated transitions (hereafter called Screen 1). Based on the responses to continuous inputs (Figure S5A-D), we focused our dynamics analysis on parameter sets that perform gene activation. Specifically, we analyzed the fraction of parameter sets that are activators and perform pulse filtering or pulse boosting (Figure 2C and Figure S5E-H separated by model family). As expected, configurations lacking TF-regulated promoter transitions are insensitive to TF dynamics (Figure 2C). Configurations that correspond to exact or approximate versions of previously established models known to perform pulse filtering or pulse boosting performed the expected function in our screen, albeit less robustly than other configurations (Figure S6A-B and Table S1). The screen indicates that configurations with a P state (3-state IPA and 4-state IPAR models) perform the most robust pulse filtering (Figure 2C). 2-state and 3-state IAR models can still perform pulse filtering but are less robust (Figure 2C and Figure S5E,G). On the other hand, only configurations with an R state (3-state IAR and 4-state IPAR models) perform pulse boosting (Figure 2C). 2- and 3-state IPA models with simple regulation are not capable of two-fold pulse boosting (Figure S5E-F), although lowering the PSR cutoff does reveal a 2-state model previously implicated in pulse boosting (Figure S6B and Table S1) (24). Interestingly, the screen also indicates that the most robust pulse boosting configurations are generally less robust than the pulse filtering configurations (Figure 2C). Together, the screen identifies the configurations that most robustly discriminate TF dynamics.

### Promoters with multi-step activation and negatively regulated deactivation robustly perform pulse filtering

#### Enriched Configuration Features

To determine properties that enable robust pulse filtering, we investigated the top 1% of configurations in Screen 1 with the highest fraction of parameter sets performing pulse filtering (Figure 3A). Each performed pulse filtering for at least 11.3% of parameter sets. Most of these configurations belong to the 3-state IPA or 4-state models, which require a transition through the intermediate P state in order to reach state A from state I. Time courses for strong pulse filtering parameter sets suggest that these configurations rely on slow priming (I→P) and/or activation (P→A) to limit promoter activation during pulses, combined with sufficiently fast deactivation to prevent accumulation of active promoter between pulses (Figure 3B-E). Additionally, all configurations have at least one negatively regulated transition out of the A state, and both TF-independent and TF-activated transcription rates are represented (Figure 3A).

**Figure 3:**
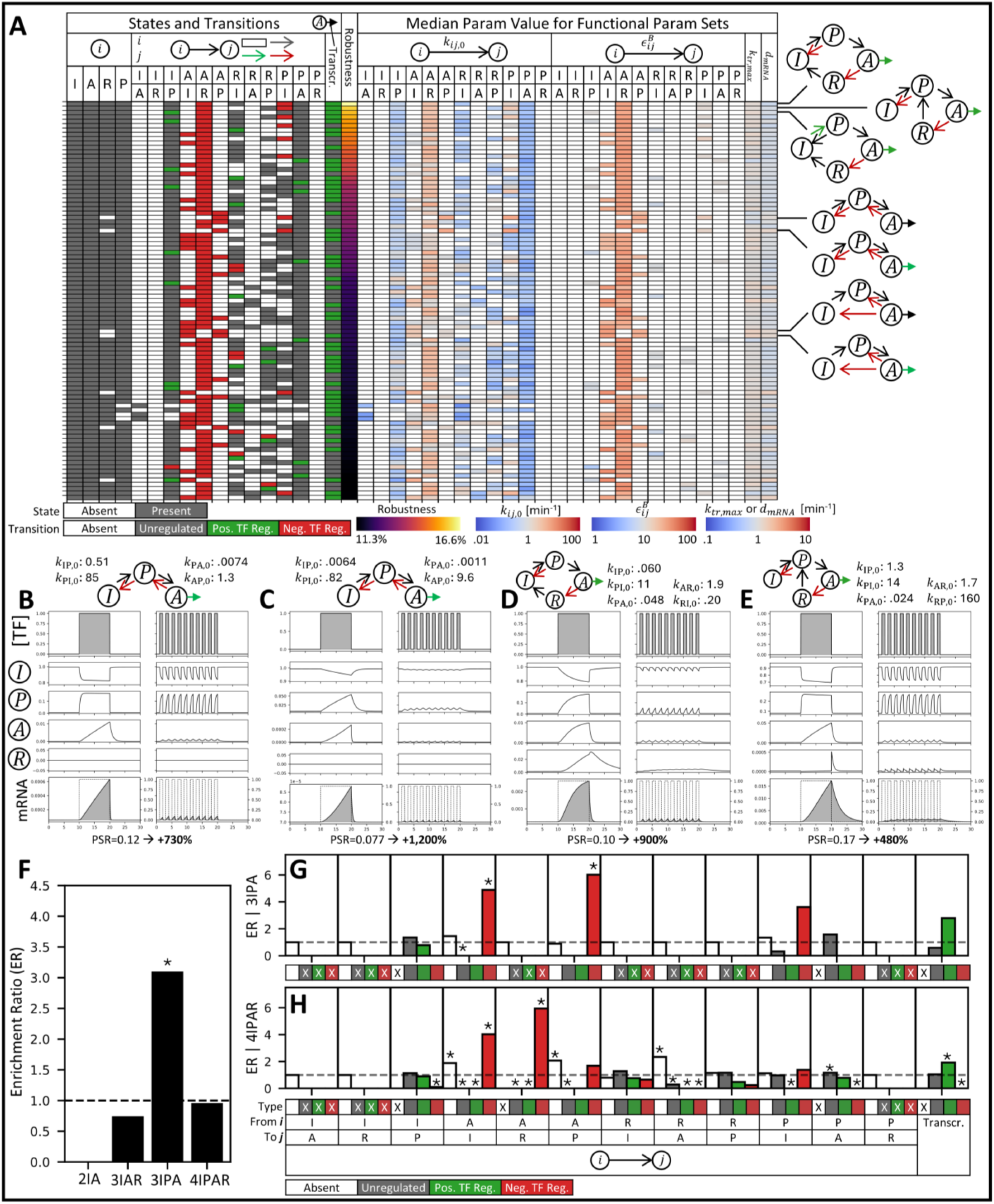
Robust pulse filtering configurations in Screen 1. A) The most robust pulse filtering configurations from Screen 1 (top 1% with the highest fraction of pulse filtering parameter sets). Each row in the table corresponds to a configuration with the columns indicating the identity of the configuration (states, transitions, transcription rate type), robustness (fraction of pulse filtering parameter sets), and parameter preferences (median parameter value across all pulse filtering parameter sets; unvaried parameters shown in white). B-E) Time courses for representative configuration/parameter sets showing the following for sustained (left) and pulsatile (right) inputs on matched axes: [TF] (x10^8^ AU_1_), promoter state fractions (I/P/A/R), and mRNA level (AU_2_) versus time (min). PSR given below. Only shown up to 10 min post-stimulation. TF-unbound rate constants (units 1/min) are shown above and all parameters are given in Supplementary Note 1. F) Enrichment ratio for each model family. G-H) Enrichment ratio for each transition type and transcription rate type, computed separately for the 3-state IPA and 4-state models. X indicates disallowed transitions. For F-H, statistically significant enrichment/depletion was determined using two-sided Fisher’s exact tests with multiple testing correction by the Benjamini-Hochberg procedure (*: adjusted p-value < 0.05; no mark: non-significant). The configurations in this figure come from the scatter plots in Figure 2C.

We quantified these observations using an enrichment ratio, which compares the frequency of a given trait among robust configurations to the expected frequency under random selection from all screened configurations (Methods). The 3-state IPA model family is significantly enriched by approximately three-fold, while the other model variants are not enriched (Figure 3F). Within 3-state IPA models, there is significant enrichment of negatively regulated A→P (∼ 6-fold) and A→I (∼5-fold) transitions (Figure 3G). Similarly, within 4-state models, there is significant enrichment of negatively regulated A→R (∼6-fold) and A→I (∼4-fold) transitions (Figure 3H). Thus, the ability of the TF to negatively regulate deactivation of the promoter is critical for robust pulse filtering.

Additionally, the 4-state models show significant enrichment for the absence of the R→A transition (Figure 3H), which implies re-activation from the R state occurs by transitioning through at least one intermediate state. Likewise, 4-state models are also enriched for the absence of the A→P and A→I transitions, which favors deactivation through the R state. Therefore, just like having a P state introduces an additional transition between the I and A states, having an R state can (in the absence of R→A) introduce an additional transition for re-activation. Consistent with this, the few 3-state IAR configurations that robustly filter pulses have a negatively regulated A→R transitions and an absent R→A transition (Figure 3A). In fact, a small number of 3-state IAR configurations are equivalent to 3-state IPA models if the states are relabeled, although this does not apply to most because the allowed transitions into/out of the P versus R states and their regulatory classifications are different.

#### Parameter Preferences

To assess parameter preferences, we computed for each configuration the median parameter values among the pulse filtering parameter sets and compared them to the middle of their respective parameter ranges (Figure 3A). All robust 3-state IPA and 4-state configurations prefer lower unbound rate constants for at least one transition along the I→P→A direction (*k*_*IP*,0_ and/or *k*_*PA*,0_), as well as higher unbound rate constants for reverse transitions out of the A state (*k*_*AP*,0_, *k*_*AI*,0_, *k*_*AR*,0_) (Figure 3A). When negatively regulated transitions are present, stronger regulation (higher 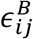) is preferred compared to for positively regulated transitions. Together, these parameter preferences align with the observations of slow priming/activation upon TF input and fast deactivation upon TF withdrawal (Figure 3B-E). Furthermore, 4-state models also show a preference for lower *k*_*RI*,0_ and/or *k*_*RP*,0_ (Figure 3A), supporting the use of the R state as an intermediate that slows re-activation and resembles priming.

#### Mechanism and Operating Regime

We next investigated the mechanism underlying robust pulse filtering and sought to understand why negatively regulated deactivation was more robust than positively regulated activation. To address this, we focused on a subset of 2-state and 3-state IPA models containing either positively regulated forward transitions (I→A or I→P→A) or negatively regulated reverse transitions (A→I or A→P→I) (Figure S7A). We note that variants of 2-state and fully reversible 3-state IPA models have been used to describe pulse filtering behavior by some Msn2-, Crz1-, and NF-κB-responsive promoters (11,12,24,26). For each configuration, we performed a gridded parameter sweep of all unbound rate constants (*k*_*ij*,0_) holding other parameters fixed (Figure 4A-B,D-E and Figure S7B-E).

**Figure 4:**
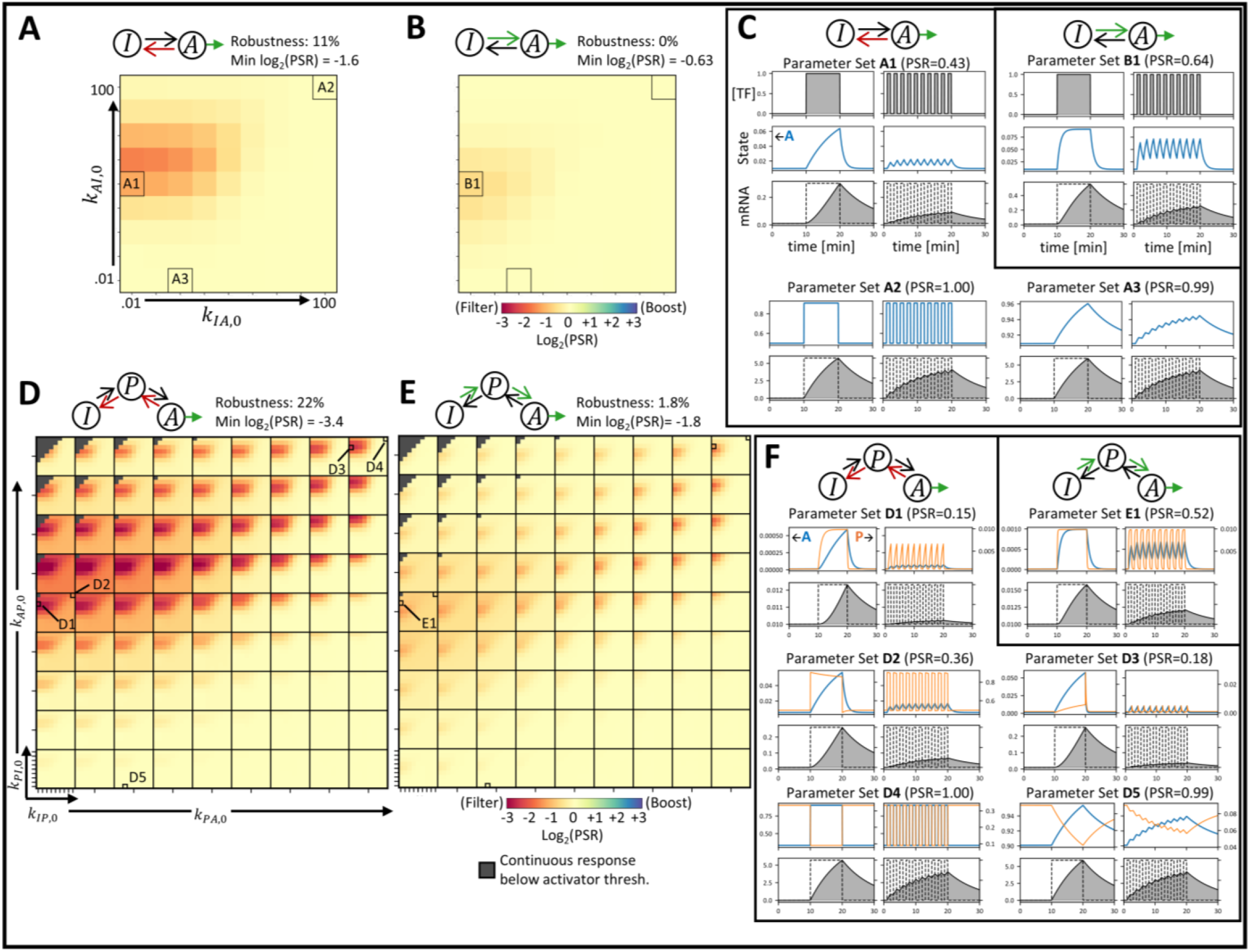
Mechanism and operating regime for pulse filtering. A,B,D,E) Heatmaps of log2 of PSR (negative/red indicates pulse filtering) for the indicated 2-state or 3-state IPA configurations showing gridded sweeps of the indicated TF-unbound rate constants (varied log uniformly from .01 to 100 min^-1^) with the base TF dynamics (Table S2: “Gridded Sweeps, Base TF Dynamics”). Non-responsive parameter sets (less than 25% increase in mRNA in response to continuous presence vs absence of TF) are masked in grey. C,F) For the indicated parameter sets from A,B,D,E time courses of the following for sustained (left) and pulsatile (right) inputs on matched axes: [TF] (x10^8^ AU_1_; shown for first parameter set only), promoter state fractions (A on left y-axis and P on right y-axis for 3-state IPA models), and mRNA level (AU_2_) versus time (min). Only shown up to 10 min post-stimulation.

In agreement with Screen 1, configurations with a P state are more robust, indicated by larger fractions of the parameter space performing pulse filtering, i.e. log_2_(PSR) < -1 (Figure 4A-B vs. 4D-E and Figure S7D-E). (Minor differences in absolute values of the robustness fractions compared to Screen 1 are due to differences in fixed and varied parameters as well as the continuous response cutoff). Also consistent with Screen 1, configurations with negatively regulated deactivation are more robust than their counterparts with positively regulated activation (Figure 4A,D vs. 4B,E and Figure S7D vs S7E). These trends hold for a wide range of TF dynamics parameters, as demonstrated for pulsatile inputs with varied *τ*_*off*_, *τ*_*on*_, or *A* compared to sustained inputs with matched AUC, amplitude, and end times (Figure S7F-G).

Although less robust, the 2-state models provide insight into mechanistic requirements for pulse filtering. First, the promoter activation timescale in presence of TF should be slow compared to *τ*_*on*_, such that the A state is not rapidly saturated during a pulse (Figure 4C; satisfied by parameter A1 but not A2). Second, the promoter deactivation timescale in absence of TF should be fast compared to *τ*_*off*_ to prevent accumulation of activated promoter between pulses (Figure 4C; satisfied by parameter A1 but not A3). These requirements resemble those previously identified for frequency modulation by a 2-state model with positively regulated activation (11). Furthermore, these requirements can explain why negatively regulated deactivation (A→I) outperforms positively regulated activation (I→A). In the presence of TF, positive regulation increases the I→A transition rate (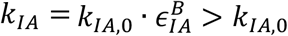 for *ρ_B_* = 1) while negative regulation decreases the A→I transition rate (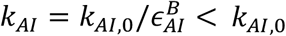 for *ρ_B_* = 1). At matched regulatory strengths 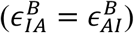, these model variants have the same steady-state levels of promoter activation in response to continuous inputs (Figure S8B-C,H-I). However, the positive regulation variant has a faster activation timescale in the presence of TF compared to negative regulation (Figure 4C parameters B1 vs A1 and Figure S8D,J), which is unfavorable for pulse filtering. Both variants have the same kinetics in the absence of TF (*k*_*IA*_ = *k*_*IA*,0_ and *k*_*AI*_ = *k*_*AI*,0_ for *ρ*_*B*_ = 0), so neither has an advantage regarding deactivation between pulses (Figure S8E,K). Together, this results in stronger pulse filtering for the negative versus positive regulation variant (Figure 4A-B and Figure S8F,L).

These mechanistic requirements can also be applied to understand the benefit of an intermediate P state (Figure 4D-E). The strongest pulse filtering occurs when both the priming/de-priming and activation/deactivation timescales satisfy the filtering requirements (Figure 4F parameter D1). However, pulse filtering can be driven by either priming/de-priming or activation/deactivation alone when the transitions between one pair of states satisfy the filtering requirements and the transitions between the other pair of states are rapid (Figure 4F parameters D2 and D3). In contrast, when the transitions are either both rapid or both slow, pulse filtering is lost (Figure 4F parameters D4 and D5). Overall, an intermediate priming state provides an additional transition in series where filtering can be achieved, increasing the maximum achievable PSR and robustness (Figure 4D-E vs Figure 4A-B).

Taken together, promoter configurations that activate through intermediate states and have deactivating transitions that are negatively regulated by the TF robustly filter pulses. These promoters perform pulse filtering in regimes where they activate slow enough to prevent saturated activation during TF pulses and deactivate fast enough to prevent accumulated activation between TF pulses.

### Promoters with TF-mediated refractoriness robustly perform pulse boosting

#### Enriched Configuration Features

We next investigated properties that enable robust pulse boosting. To do so, we investigated the top 1% of configurations in Screen 1 with the highest fraction of parameter sets performing pulse boosting (Figure 5A). These configurations have at least 2.2% of parameter sets performing pulse boosting (compared to 11.3% for the top pulse filtering ones), indicating that promoter configurations with simple regulation are less capable of robust pulse boosting than pulse filtering. These configurations all have an R state, with the 3-state IAR model being significantly enriched about four-fold (Figure 5A,F). In all configurations, the TF negatively regulates at least one transition out of the R state, which serves to trap the promoter in the R state when bound by the TF (Figure 5A). Indeed, negatively regulated R→I and R→A transitions are significantly enriched more than three-fold within 3-state IAR models (Figure 5G), and negatively regulated R→I, R→A, and R→P transitions are all significantly enriched more than two-fold within 4-state models (Figure 5H). Within 4-state models these transitions are also significantly enriched for absence, indicating they should be negatively regulated or not permitted. Time courses of strong pulse boosting show that the R state restricts promoter activation at longer durations, in some cases following an initial overshoot in promoter activation (Figure 5B-E). For sustained inputs, refractoriness persists for much of the stimulation window. For pulsatile inputs, a full or partial recovery between pulses allows repeated activation and transcription at the beginning of each pulse. Additionally, every configuration has a TF-activated transcription rate (Figure 5A,G-H), which is required for steady-state activation of mRNA transcription in the presence of refractory-associated promoter deactivation.

**Figure 5:**
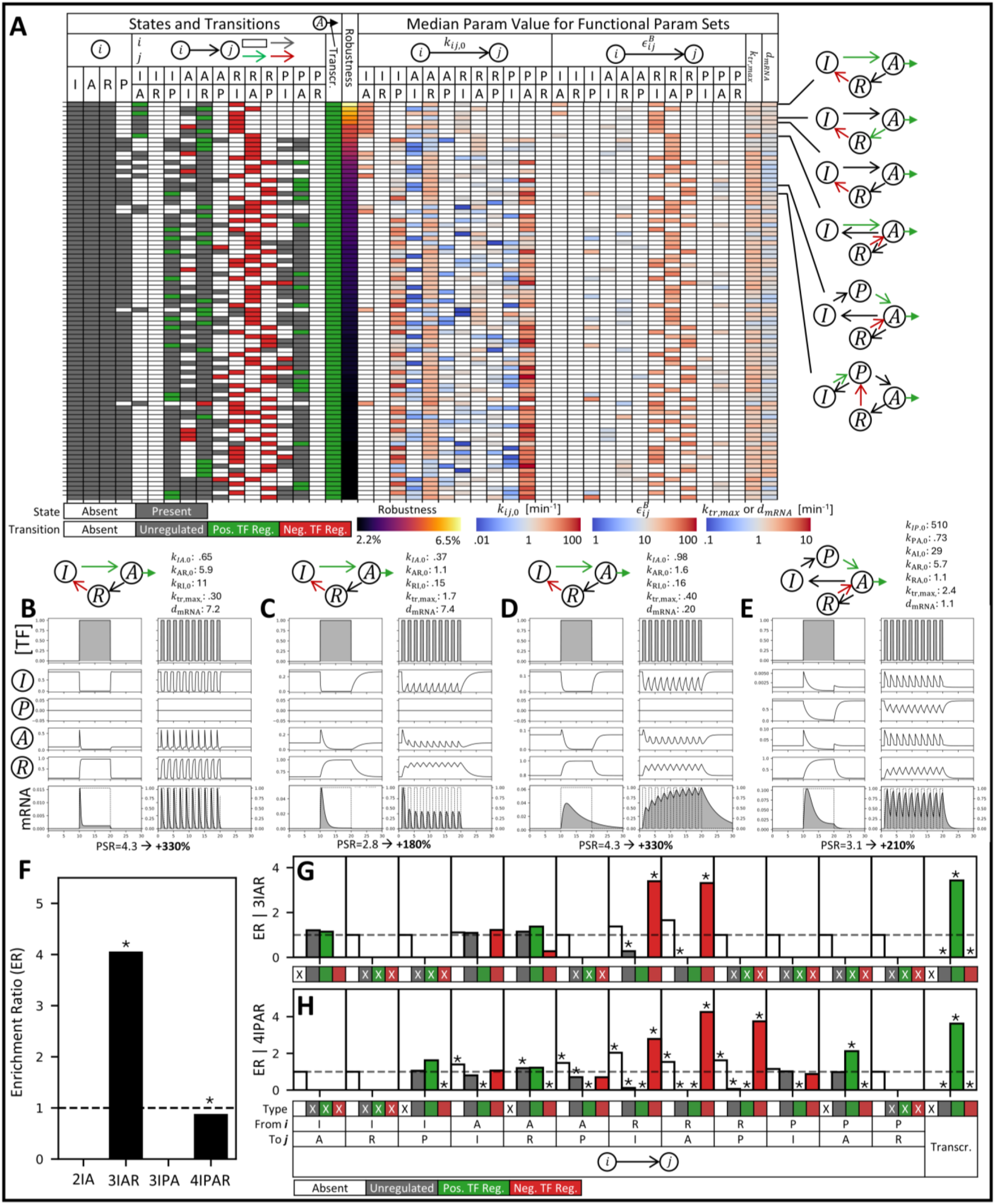
Robust pulse boosting configurations in Screen 1. A) The most robust pulse boosting configurations from Screen 1 (top 1% with the highest fraction of pulse boosting parameter sets). Each row in the table corresponds to a configuration with the columns indicating the identity of the configuration (states, transitions, transcription rate type), robustness (fraction of pulse boosting parameter sets), and parameter preferences (median parameter value across all pulse boosting parameter sets; unvaried parameters shown in white). B-E) Time courses for representative configuration/parameter sets showing the following for sustained (left) and pulsatile (right) inputs on matched axes: [TF] (x10^8^ AU_1_), promoter state fractions (I/P/A/R), and mRNA level (AU_2_) versus time (min). PSR given below. Only shown up to 10 min post-stimulation. Rate constants (units 1/min) are shown above and all parameters are given in Supplementary Note 1. F) Enrichment ratio for each model family. G-H) Enrichment ratio for each transition type and transcription rate type, computed separately for the 3-state IAR and 4-state models. X indicates disallowed transitions. For F-H, statistically significant enrichment/depletion was determined using two-sided Fisher’s exact tests with multiple testing correction by the Benjamini-Hochberg procedure (*: adjusted p-value < 0.05; no mark: non-significant). The configurations in this figure come from the scatter plots in Figure 2C.

#### Parameter Preferences

Most configurations favor fast activation kinetics, as demonstrated by the higher *k*_*IA*,0_ values for 3-state IAR models and higher *k*_*IP*,0_ and *k*_*PA*,0_ values for 4-state models (Figure 5A). This ensures sufficient promoter activation at the onset of TF stimulation (during both sustained and pulsatile inputs), which contrasts with the parameter preferences for pulse filtering. Although less pronounced, a fast transition into the R state also appears favored, as seen by high *k*_*AR*,0_ values for many configurations, especially those where the A→R transition is not already accelerated by positive regulation of this transition (Figure 5A). Trends for the unbound rate constants were not clear for the exit from the R state, but strong regulatory strengths 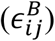 are preferred for these transitions (Figure 5A).

#### Mechanism and Operating Regime

To better understand the mechanism behind robust pulse boosting, we focused on versions of the 3-state IAR model with regulatory features identified in Screen 1: either positively regulated transitions into the R state or negatively regulated transitions out of the R state (Figure S9A). We also considered a 2-state model with TF-independent transcription rate (Figure S10A), because it showed weak pulse boosting in Screen 1 (Figure S6B) and resembles a model implicated in pulse boosting by some Crz1-responsive promoters (24). For each, we performed a gridded parameter sweep of all unbound rate constants (*k*_*ij*,0_) holding other parameters fixed (Table S2: “Gridded Sweep, Base Dynamics”). In agreement with Screen 1, the 3-state IAR models with negatively regulated transitions out of the R state (Figure 6A and Figure S9B) are more robust at pulse boosting than their counterparts with positively regulated transitions into the R state (Figure 6B and Figure S9C), as well as compared to the 2-state model (Figure S10B). This trend holds for a wide range of TF dynamics parameters, as demonstrated for pulsatile inputs with varied *τ*_*off*_, *τ*_*on*_, or *A* compared to sustained inputs with matched AUC, amplitude, and end times (Figure S9D-E).

**Figure 6:**
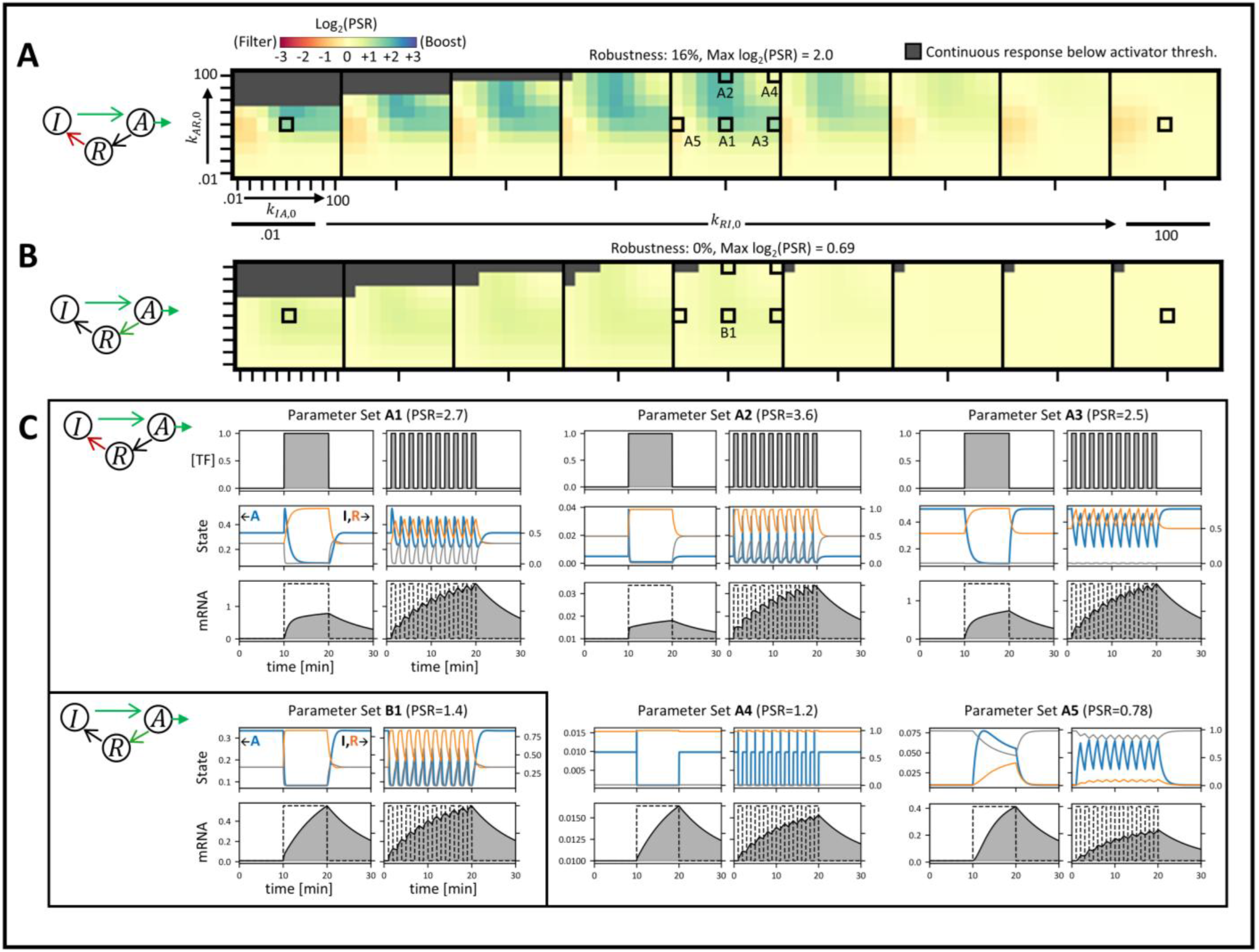
Mechanism and operating regime for pulse boosting. A-B) Heatmaps of log2 of PSR (positive/blue indicates pulse boosting) for the indicated 3-state IAR configurations showing gridded sweeps of the indicated TF-unbound rate constants (varied log uniformly from .01 to 100 min^-1^) with the base TF dynamics (Table S2: “Gridded Sweeps, Base TF Dynamics”). Non-responsive parameter sets (less than 25% increase in mRNA in response to continuous presence vs absence of TF) are masked in grey. C) For the indicated parameter sets from A-B time courses of the following for sustained (left) and pulsatile (right) inputs on matched axes: [TF] (x10^8^ AU_1_; shown for first parameter set only), promoter state fractions (A on left y-axis and I and R on right y-axis), and mRNA level (AU_2_) versus time (min). Only shown up to 10 min post-stimulation.

Analysis of the 2-state model confirms its ability to perform weak pulse boosting (max PSR ∼ 1.5) and identifies its mechanistic requirements (Figure S10). Pulse boosting in this model relies on sustained promoter activation between pulses and thereby, due to the TF-independent transcription rate, continued transcription during the off pulse phase. This requires that activation be fast relative to *τ*_*on*_ to ensure sufficient activation during each pulse, and that deactivation be slow relative to *τ*_*off*_ to keep the promoter active and transcribing between pulses (Figure S10B-C; satisfied by parameter B1 but not B2 or B3). This provides a mechanism for weak pulse boosting that does not rely on refractoriness.

We next investigated how the 3-state IAR models use refractoriness to perform strong pulse boosting (Figure 6A-B). Following TF input, the pulse boosting parameter sets show a rise in the R state and an associated decrease in the A state, which is equally or more pronounced in sustained versus pulsatile inputs and is recovered partially or fully between pulses (Figure 6C parameters A1, A2, A3). Depending on the configuration and regulation strengths, promoter deactivation may be preceded by a transient overshoot in promoter activation. As a result of differential deactivation, the time-weighted level of A state is higher for pulsatile than sustained inputs, leading to pulse boosting at the mRNA level. For optimal pulse boosting, the decrease in A state should be fast enough to reach a minimum during the sustained input but slow enough to consume a considerable portion of a single pulse, thereby raising the time-weighted A state fraction (Figure 6C parameters A1, A2, A3). In support of this, if the decrease in A state upon TF input is rapid compared to *τ*_*on*_, pulse boosting performance is worsened (Figure 6C parameters A4). Also, when no reduction in A state is realized over the duration of the sustained input, configurations are insensitive to dynamics or even perform weak pulse filtering in the presence of a slow, transient overshoot in promoter activation (Figure 6C parameter A5). In comparison to the models with negatively regulated R→I, the models with positively regulated A→R are less efficient at pulse boosting because they show accelerated promoter deactivation upon TF input (Figure 6C parameters A1 versus B1).

Taken together, the most robust pulse boosting configurations possess a refractory state with exit transitions negatively regulated by the TF. This TF-mediated refractoriness causes promoter deactivation that saturates for sustained inputs and recovers for pulsatile inputs, thereby enabling pulse boosting.

### Few configurations can perform both pulse filtering and boosting

We also searched for configurations in Screen 1 that could perform pulse boosting under some parameter sets and pulse filtering under others. Only 5 of the 8,828 configurations have >2.5% parameter sets classified as pulse boosting and >2.5% classified as pulse filtering (Figure S11). These configurations all possess an R state and two regulated transitions: a negatively regulated transition out of the A state, which is implicated in pulse filtering, and a negatively regulated transition out of the R state, which is implicated in pulse boosting (Figure S11A). As expected, the parameter preferences for pulse filtering versus pulse boosting are often opposite, e.g. the unbound rate constants for the I→A or P→A transitions are lower for pulse filtering parameter sets and higher for pulse boosting parameter sets (Figure S11C-D).

### Promoters regulated by bifunctional TFs extend the design space for pulse boosting

Most natural TFs are thought to function exclusively as activators or repressors. Some TFs, however, can activate or repress depending on contextual factors including cell type, chromatin state, and availability of cofactors (38,39). The majority likely act as activators of certain genes and repressors of others. However, TFs bearing both activator and repressor domains or effector domains capable of both activation and repression exist (40,41), which may enable these TFs to activate and repress the transcription of a single target gene within a single context. Indeed, a recent study focused on this type of TFs exerting so-called incoherent regulation under a fixed molecular context (42). Moreover, synthetic TFs harboring combinations of activator and repressor domains have been recently engineered (43), suggesting that this type of regulation is also becoming available in synthetic biology.

To assess the capabilities of bifunctional TFs, we focused on a second set of configurations. Configuration set 2 includes all 2- and 3-state configurations including both simple and bifunctional TF-mediated regulation and with no limit on the total number of regulated transitions (Figure 1G and Figure S1G). As our analysis focuses on the parameter sets that behave as activators in response to continuous TF input, which was not achievable by any configurations with a TF-repressed transcription rate in screen 1 (Figure S5A-D), we only analyzed configurations with TF-independent and TF-activated transcription rates, leaving 2,070 of the 3,105 configurations from configuration set 2 (hereafter referred to as Screen 2).

In Screen 2, the most robust pulse filtering is achieved exclusively by configurations with simple TF regulation (Figure 7A,E). In agreement with Screen 1, these configurations belong mostly to the 3-state IPA models (19/20 of the top 1%) and possess at least two negatively regulated transitions out of state A (Figure 7C). In contrast, the most robust pulse boosting in Screen 2 is achieved exclusively by configurations with bifunctional TF regulation (Figure 7B,G). While many still belong to the 3-state IAR model and possess TF-regulated transitions that promote trapping in the R state (i.e. negatively regulated R→I/A and/or positively regulated A→R), multiple 3-state IPA models and one 2-state model appear in the top 1% most robust pulse boosting configurations (Figure 7D). These configurations possess negatively regulated transitions into state A (I→A or P→A) as well as positively regulated transitions out of state A (Figure 7D). Therefore, these configurations combine repressor-like regulation of promoter state transitions (TF-based repression of the A state) with activator-like regulation of transcription from the A state (TF-activated transcription rate). Time courses show that TF input leads to an initial phase of elevated transcription followed by promoter deactivation at longer times, thereby favoring multiple shorter pulses over a single longer one (Figure S12B-D). This mechanism directly resembles the refractory-based mechanism described in Screen 1 and is capable of even stronger pulse boosting. Lastly, Screen 2 also indicates differences in the regulatory complexity required for optimal pulse filtering versus boosting. For pulse filtering, the ideal number of regulated transitions is 40-80% of the total number of transitions (Figure 7F), while for pulse boosting, regulation of 80-100% of the total number of transitions is ideal (Figure 7H).

**Figure 7:**
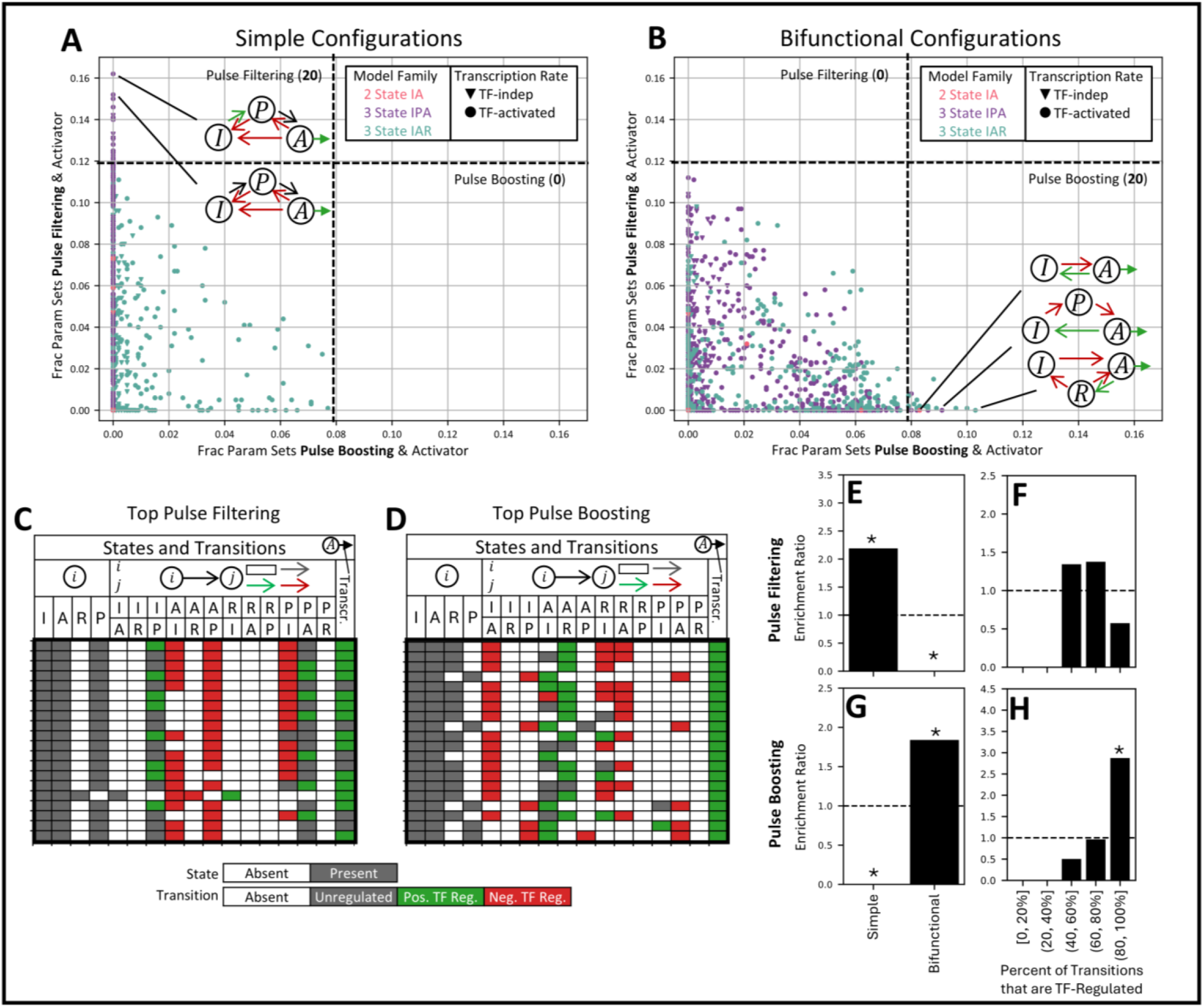
Bifunctional TF regulation extends the design space for pulse boosting. A-B) Response of each configuration in screen 2 to TF dynamics, separated by Simple (A) and Bifunctional (B) configurations. Scatter plots show the fraction of parameter sets that are activators and perform pulse filtering (PSR<1/2) versus pulse boosting (PSR>2). Dotted lines correspond to cutoffs containing the top 1% most robust pulse filtering and pulse boosting configurations across the entire screen 2. C-D) Top 1% most robust pulse filtering (C) and pulse boosting configurations. E-F) Enrichment ratio for simple vs bifunctional configurations (E) and percentage of transitions that are TF-regulated (F) for the top 1% most robust pulse filtering configurations. G-H) Same for the top 1% most robust pulse boosting configurations. For E-H, statistically significant enrichment/depletion was determined using two-sided Fisher’s exact tests with multiple testing correction by the Benjamini-Hochberg procedure (*: adjusted p-value < 0.05; no mark: non-significant).

## Discussion

It has been established that eukaryotic promoters can decode TF dynamics, resulting in distinct gene expression responses (1,11,12,23–27,29,32). Identifying the general design principles governing decoding at the promoter level would advance our understanding of natural gene regulation, including the promoter architectural features and TF properties required. This knowledge would also enable the engineering of compact synthetic circuits that process temporal TF signals at the promoter level.

To this aim, we used mathematical modeling to systematically elucidate the properties of promoters and TF-based regulation that enable robust decoding of TF dynamics. We found that configurations where the promoter activates via intermediate, non-transcribing states and in which bound TF negatively regulates deactivation (versus positively regulating activation) are the most robust at pulse filtering. These promoters operate in regimes where they activate slowly enough to not saturate during a TF pulse and deactivate quickly enough to prevent accumulated activation between TF pulses. Configurations with a refractory state and in which bound TF regulates occupancy of this state, especially by negatively regulating transitions out of it, are the most robust at pulse boosting. These promoters operate in regimes where a transient phase of high transcription (sometimes accompanied by an overshoot in promoter activation) is followed by refractoriness at longer timescales. Bifunctional TFs that exert activator- and repressor-like regulation extend the design space for robust pulse boosting and mechanistically resemble TF-mediated refractoriness (which we considered neither activator-nor repressor-like given limited knowledge on how TFs regulate refractoriness).

The mechanistic basis of promoter priming is relatively well characterized. Mechanisms include DNA looping, which may in some cases be mediated by the TF itself, as well as chromatin remodeling processes such as nucleosome repositioning, deposition, or eviction, which require ATP-dependent remodeling complexes (12,23,28,32). In these contexts, TFs either directly alter DNA conformation or recruit cofactors that modify chromatin accessibility and histone marks. The multiple intermediate states and slower timescales associated with histone modifications and chromatin remodeling may be especially conducive to robust pulse filtering.

By contrast, the mechanistic basis of promoter refractoriness remains to be fully understood. It is well established that eukaryotic genes are transcribed in bursts separated by inactive intervals during which the promoter is refractory to further stimulation (28,30,33–35). Refractoriness varies between genes (30), and has been linked to properties including the core promoter, cis-regulatory elements, and chromatin remodeling, depending on context (33–35). Notably, TFs still bind refractory promoters (33), and mathematical models in which the TF regulates promoter transitions associated with the R state have been able to describe experimental data (28). However, direct evidence of TFs regulating these promoter transitions is lacking. Disentangling TF-driven effects from chromatin resetting and other processes remains experimentally challenging, particularly in real-time, single-cell settings.

Our work also suggests that whether TFs regulate transitions into versus out of key promoter states has implications for robustness decoding. For instance, negatively regulating promoter deactivation is a more robust design for pulse filtering than positively regulating activation, even when both result in the same steady-state response to continuous stimulation. Interestingly, these two forms of regulation have been linked to different aspects of transcriptional bursting. In the two-state telegraph model, TF-regulated activation (I→A) modulates burst frequency while TF-regulated deactivation (A→I) modulates burst duration (28). Bursting parameters are controlled by factors including local chromatin environment, histone modifications, DNA looping, number and affinity of cis-regulatory elements, and TF availability (31,34,44–46). In some cases, bursting parameters may be uncoupled and involve different molecular mechanisms. For instance, some mechanisms have been shown to mainly control burst frequency, while others mainly control burst size (related to burst duration) (34,44–46). This suggests that different mechanisms could underly the regulation of promoter activation kinetics versus deactivation kinetics, which would have implications for understanding decoding in natural systems and implementing decoding in synthetic systems. Methods for inferring promoter state dynamics from single-cell gene expression trajectories (27), combined with stochastic modeling of gene expression, will be important in understanding how TFs regulate promoter state dynamics.

Our results further indicate that bifunctional TFs expand the space of configurations capable of robust pulse boosting. Such TFs exert opposing regulatory effects on the same promoter, such as increasing the rate of transcription from the active promoter while simultaneously inducing promoter deactivation. Although bifunctional TFs with both activator and repressor functions have been described (38–42), they are less common than TFs functioning exclusively as activators or repressors. This asymmetry may help explain why pulse filtering appears more prevalent than pulse boosting among natural promoters.

In contrast to decoding TF dynamics at the level of the target promoter, temporal signals can also be decoded at the level of downstream regulatory networks (13–17). These network motifs often make use of branches driven by different biochemical processes and occurring at different timescales and/or cellular compartments (1,16). For example, coherent feed forward loops (FFLs) with AND logic robustly perform temporal filtering by using direct and indirect branches to delay the on response time (15,16). Conversely, incoherent FFLs have been shown to selectively respond to pulsatile versus sustained inputs using separate activation and inhibition branches (14,15,17). In contrast, our framework consists of a single target gene regulated by a single input (the TF), without feedback regulation from the target gene to the TF or feed-forward regulation involving cofactors (which implicitly are assumed to have constant availability). Future work should address advantages and disadvantages of promoter-vs network-level decoding in terms of both theoretical performance and biological complexity.

Pulse filtering has an intuitive biological rationale: it prevents inappropriate gene activation in response to transient or noisy stimuli and avoids premature commitment to irreversible outcomes such as apoptosis. The physiological relevance of pulse boosting, by contrast, is conceptually less straightforward, but it may be advantageous in contexts such as circadian regulation or repeated environmental stimuli. Nevertheless, in synthetic systems, the ability to differentially regulate gene expression programs by externally controlling TF dynamics would offer powerful regulatory flexibility. Advances in engineering controllable bifunctional TFs may therefore enable dynamic-input-based gene regulation without the need for complex circuits.

### Limitations of the study

We considered promoter configurations with one active state and at most one primed and one refractory state, which allowed us to include a range of models previously used to describe the sensitivity of promoters to TF dynamics (11,12,23–26,28). The framework presented here could be extended to more complex models with multiple primed (32), refractory (30), and/or active (27) states. Secondly, our study focused on decoding in the context of gene activation. Future work should explore additional regimes, including temporal selectivity in gene repression, as well as systems exhibiting steady-state repression with transient activation. Thirdly, we assessed the total transcriptional output in response to different inputs. As the dynamics of gene expression itself can also influence cell responses, future work could add this to the functional characterization. For instance, p53 target genes exhibit different expression dynamics depending on the relationship between pulse frequency and mRNA and/or protein stability (47). Fourthly, our screens were performed under a saturating TF condition, and we therefore did not explore preferences for parameters controlling the equilibrium bound fraction (*K*_*d,TF*_ and *n*_*TF*_). Lastly, we used a deterministic framework. Stochastic modeling could provide insight into how noise interacts with temporal selectivity and could let us explore connections between bursting parameters and TF-based regulation of promoter dynamics.

## Methods

### Model Formulation

The model of TF-regulated gene expression considers a sequence-specific TF that binds to the enhancer (one or more response elements) upstream of the core promoter driving the target gene. TF:DNA binding/unbinding is assumed to be sufficiently fast compared to other processes (including promoter state transitions), such that the bound fraction is at equilibrium, as commonly assumed (28). The equilibrium bound fraction (*ρ*_*B*_) is modeled as a function of the nuclear concentration of active TF, [TF], using a Hill-type function to account for potential cooperative TF binding:

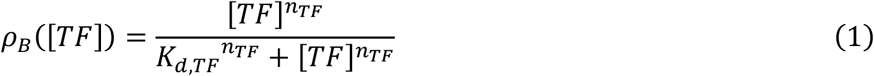

The promoter exists in one of multiple functional states *P*_*s*_ where *s* ∈ {*I, P, A, R*} and transitions between these states with rate constants that account for contributions from the equilibrium fraction in the bound and unbound states. The rate constant for the transition of the promoter from state *P*_*i*_ to *P*_*j*_ is then defined as:

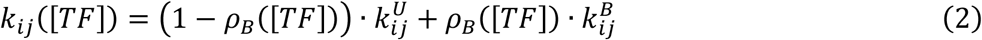

where 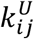 and 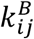 are the rate constants for the transition from state *P*_*i*_ to *P*_*j*_ for the unbound and bound promoter, respectively. We assume that the binding/unbinding parameters are the same for all promoter states.

Transitions between states can be regulated by bound TF. A formalism for TF-regulated promoter state transitions from Li et al. (28) is used in which a transition can be positively regulated/accelerated by a bound TF, negatively regulated/decelerated by a bound TF, or independent of the TF. Thus, the transitions *T*_*ij*_ from promoter state *P*_*i*_ to *P*_*j*_ takes one of the following four types:

*T*_*ij*_ = 0: absent (there is no transition)

*T*_*ij*_ = 1: TF-independent (bound TF does not affect the rate)

*T*_*ij*_ = 2: positively regulated by the TF (bound TF increases the rate)

*T*_*ij*_ = 3: negatively regulated by the TF (bound TF decreases the rate)

The rate constant for the transition of the promoter from state *P*_*i*_ to *P*_*j*_ depends on the form of regulation as follows:

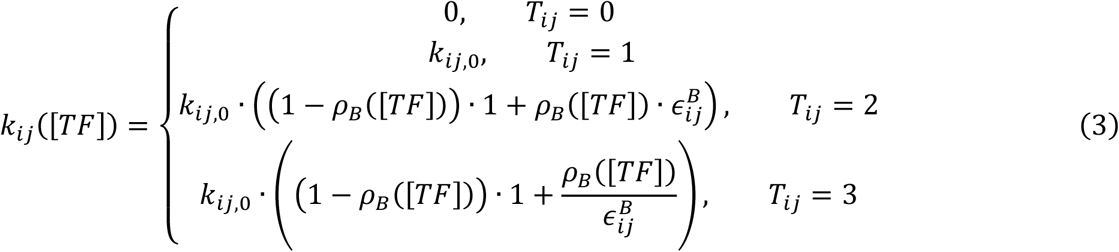

where *k*_*ij*,0_ is the TF-unbound rate constant for the transition of the promoter from state *P*_*i*_ to *P*_*j*_ and 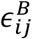 is the strength of TF regulation for this transition. Note that 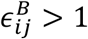.

Transcription of mRNA occurs from the *P*_*A*_ state. Two commonly used forms for the rate of transcription were considered: TF-independent, where transcription from the A state is constant regardless of bound TF (24,25,28), and TF-dependent, where transcription from the A state depends on the fraction of bound TF (12,23,28), being either TF-activated or TF-repressed. The transcription rate is given by:

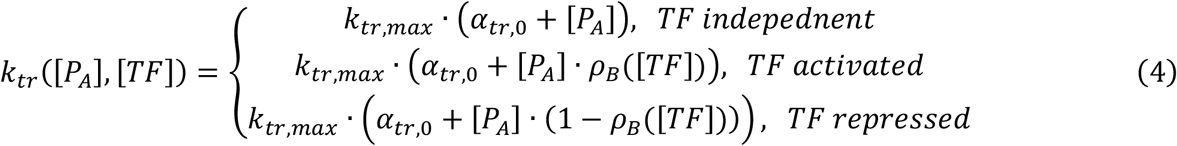

where *k*_*tr,max*_ is the maximum regulated transcription rate and *α*_*tr*,0_ is the relative basal transcription constant.

The population-level dynamics of the model are described using a system of ordinary differential equations (ODEs), as done previously for models of TF-regulated transcription from multi-state promoters (11,12,23–25,28). The system is comprised of ODEs for the fraction of promoters in each of four states *P*_*i*_ and for the abundance of mRNA, as well as the conservation of total promoter fraction:

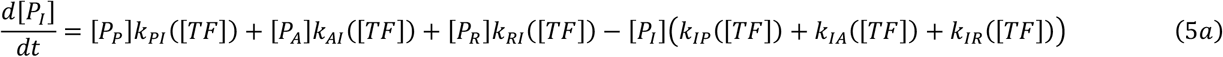

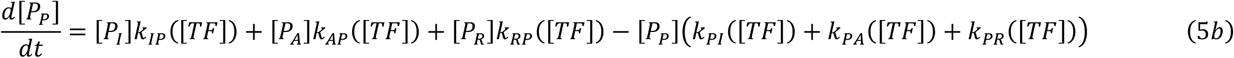

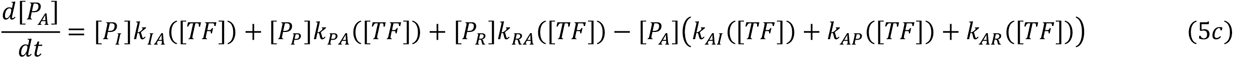

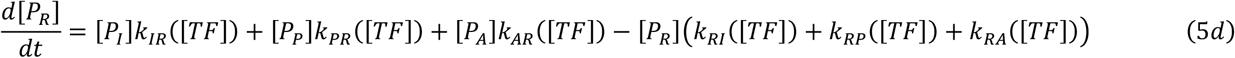

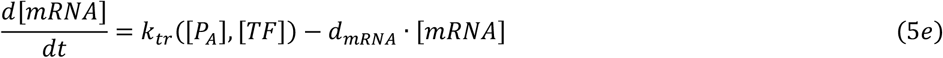

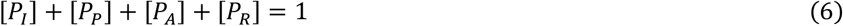

where *k*_*ji*_([*TF*]) and *k*_*tr*_([*P*_*A*_], [*TF*]) have already been defined above and *d*_*mRNA*_ is the mRNA degradation rate. The 2- and 3-state promoter models can be described using this generalized 4-state system.

### TF Dynamics

The input to the model is a time-dependent nuclear concentration of active TF (“TF dynamics”). We considered two TF dynamics: sustained (a single long pulse) and pulsatile (multiple shorter pulses). These are motivated by the dynamics of natural TFs and previous experimental and theoretical work on the sensitivity of genes to TF dynamics (1,2,12,23,24). The pulsatile input is a square wave parameterized by the pulse on time *τ*_*on*_, pulse off time *τ*_*off*_, number of pulses *N*_*p*_, and amplitude *A*:

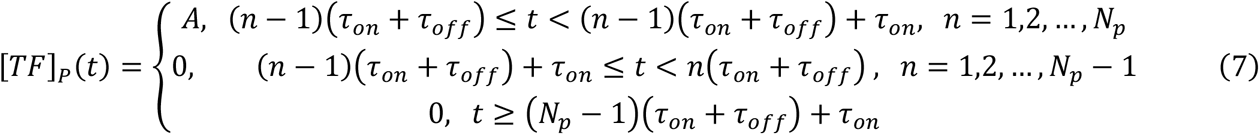

Note that an alternative parameterization of the pulsatile TF input dynamics using frequency 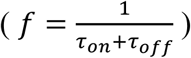 and duty cycle 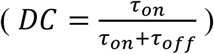 is also possible.

The pulsatile input was compared to a sustained input with a matched [TF] area under the curve (AUC) and amplitude, which was previously done to exclude effects due to cumulative TF levels (4,23). For a pulsatile input parameterized by pulse on time *τ*_*on*_, pulse off time *τ*_*off*_, number of pulses *N*_*p*_, and amplitude *A*, the corresponding AUC- and amplitude-matched sustained input consists of a single pulse with on time *N*_*p*_*τ*_*on*_ and amplitude *A*, yielding the following equation:

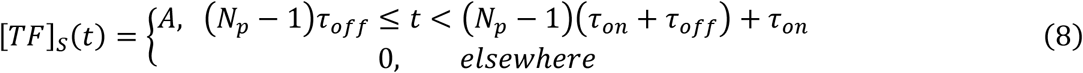

where the end points of the stimulation phases for the pulsatile and sustained inputs are also aligned, as previously done (23).

In addition to dynamic inputs, the steady state response to continuous absence and presence of TF was also analyzed using the following inputs: [*TF*]_*Continous Off*_(*t*) = 0 and [*TF*]_*Continous On*_(*t*) = *A*.

### Configuration Enumeration

A promoter configuration is uniquely defined by its states, transition types, and transcription rate type. The states are given by the vector *S*_*i*_ = [*S*_*I*_, *S*_*A*_, *S*_*R*_, *S*_*P*_] where *S*_*i*_ ∈ {0,1} indicates the absence (0) or presence (1) of each state. The transition types are given by the matrix *T*_*ij*_ where *T*_*ij*_ ∈ {0,1,2,3} indicates the transition type for the transition from state *i* to state *j*, which is either absent (0), TF-independent (1), positively regulated by the TF (2), or negatively regulated by the TF (3), as defined above. The transcription rate type is given by *V*_*tr*_ where *V*_*tr*_ ∈ {1,2,3} indicate TF-independent (1), TF-activated (2), and TF-repressed (3) transcription rates, as defined above.

We enumerated all 2-, 3-, and 4-state promoter configurations that satisfied the following rules. 2-state IA configurations consist of states I and A. 3-state IPA configurations consist of states I, P, and A. 3-state IAR configurations consist of states I, A, and R. 4-state configurations consist of states I, P, A, and R. Regarding state connectivity, every state must be reached from every other state via a sequence of allowable state transitions. Transitions from state I to state R or state P to state R are not allowed because state R is assumed to only be reached from state A (i.e. after activation). If state P is present, the transition from state I directly to state A is not allowed, because priming is assumed to be required for activation.

### Configuration Classification

Configurations were classified according to three properties:

1. States present, i.e. 2-State IA, 3-State IPA, 3-State IAR, or 4-State IPAR model families.
2. Number of TF-regulated transitions.
3. Simple vs Bifunctional TF-based regulation. Simple regulation requires that the TF does not exert both activator-like and repressor-like regulation, while for bifunctional regulation the TF does exert both. The following were defined as activator-like: positive regulation of transitions on the path from I to A (I→A, I→P, P→A), negative regulation of the corresponding reverse transitions (I←A, I←P, P←A), and TF-activated transcription rate. Conversely, the following were defined as repressor-like: negative regulation of forward transitions along the path from I to A (I→A, I→P, P→A), positive regulation of the corresponding reverse transitions (I←A, I←P, P←A), and TF-repressed transcription rate. As there is incomplete evidence of how TFs influence transitions to/from the R state, all transitions to/from the R state were considered unclassified, i.e. neither activator-nor repressor-like.

### Configuration Sets

As it was not feasible to screen all the over 324,000 enumerated configurations with sufficient parameter space coverage, two configuration subsets were investigated. Configuration Set 1 contains all 2-, 3-, and 4-state configurations with a maximum of 2 TF-regulated promoter transitions and in which the TF exerts simple regulation (8,828 configurations in total). To investigate behaviors resulting from bifunctional regulation and higher numbers of TF-regulated promoter transitions, Configuration Set 2 consists of all 2- and 3-state configurations, with no restrictions on the number or type of regulation exerted by the TF (3,105 configurations in total, of which 625 overlap with Configuration Set 1).

### Configuration Analysis

To screen each configuration for the ability to robustly discriminate TF dynamics, Latin Hypercube Sampling (LHS) (48) was used to generate 1,000 parameter sets according to the parameter ranges and distributions specified in Table S2. As the relative results were similar using a non-saturating or saturating TF concentration, Screens 1 and 2 were performed with a saturating TF concentration (Table S2), which corresponds to the TF-bound promoter fraction being approximately 1 in the presence of TF and 0 in the absence of TF.

For each parameter set, the steady-state mRNA levels in response to a continuous presence ([*TF*] = *A*) and absence ([*TF*] = 0) of TF were first assessed. The fold change in steady-state mRNA levels was defined as:

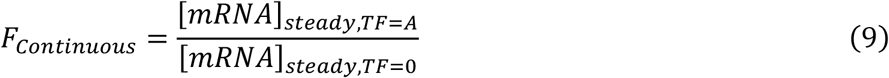

Using this metric, parameter sets were classified as steady state activators (*F*_*Continuous*_ ≥ 2), repressors (*F*_*Continuous*_ ≤ 1/2), or non-responsive (1/2 < *F*_*Continuous*_ < 2).

To assess sensitivity to TF dynamics, we analyzed gene expression in response to sustained versus pulsatile TF inputs (defined above). The sustained and pulsatile inputs were set to end at the same timepoint, have the same post-stimulation wait duration (60 min), and have the same total simulation duration. Numerical simulations were conducted with the initial condition set to the steady-state promoter state fractions and mRNA levels in the absence of TF ([*TF*] = 0). Responses to sustained and pulsatile inputs were computed using the mRNA AUC. Specifically, for sustained and pulsatile inputs the AUC of mRNA over the simulation referenced to the baseline mRNA level (i.e. in absence of TF) was computed:

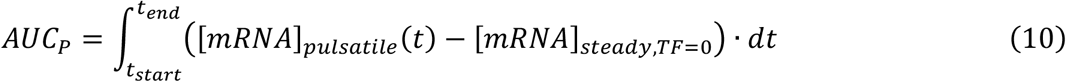

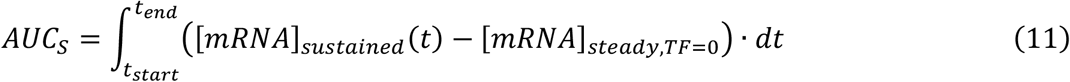

To compare the response to pulsatile versus sustained inputs, the ratio of the mRNA AUC for the pulsatile to sustained inputs was used, referred to as the Pulsatile-to-Sustained Ratio (PSR):

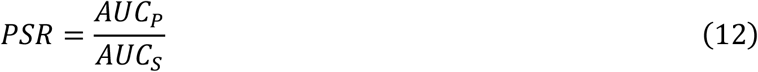

Using the *PSR*, parameter sets were classified as pulse filtering (*PSR* ≤ 1/*C*), pulse boosting (*PSR* ≥ *C*), or insensitive to TF dynamics. For Screens 1 and 2, the cutoff *C* = 2 was used. Note that only parameter sets that responded as activators to continuous inputs were considered, all of which had non-negative values for both *AUC*_*P*_ and *AUC*_*S*_. In computing the *PSR*, if both *AUC*_*P*_ and *AUC*_*S*_ were not at least 0.1% larger than the baseline, 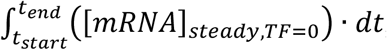, they were considered non-significant and the *PSR* was set to 1 (i.e. insensitive). If only one AUC was non-significant, then it was set to 0.1% above the baseline for computing the *PSR*.

### Configuration-level Metrics

In computational screens that evaluate the robustness of biological systems to parameter variations, the robustness is often defined as the fraction of parameter sets for which a target function is performed above a threshold score (13,14,36). Therefore, for each promoter configuration we computed the pulse filtering robustness as the fraction of parameter sets that perform activation and pulse filtering (PSR < 1/2), and the pulse boosting robustness as the fraction of parameter sets that perform activation and pulse boosting (PSR > 2). As we focus on gene activation, this definition reflects that only parameter sets that responded as activators to continuous inputs were considered in the dynamic response analysis.

To determine properties that enable robust pulse filtering or pulse boosting, we investigated the top ∼1% of configurations with the highest pulse filtering robustness or pulse boosting robustness, respectively. For Screen 1 pulse filtering, this corresponded to the top 91 configurations (each having at least 11.3% of parameter sets that perform pulse filtering). For Screen 1 pulse boosting, this corresponded to the top 89 configurations (each having at least 2.2% of parameter sets that perform pulse boosting). For Screen 2 pulse filtering, this corresponded to the top 20 configurations (each having at least 12.1% of parameter sets that perform pulse filtering). For Screen 2 pulse boosting, this corresponded to the top 20 configurations (each having at least 8.0% of parameter sets that perform pulse boosting). As an indicator of the parameter preferences of robust pulse filtering/boosting for a given configuration, the median parameter value for the pulse filtering/boosting parameter sets was used. We note that the median parameter value cannot differentiate between a lack of preference and a preference for the middle of the parameter range. The robust configurations were displayed using colored tables where each row corresponds to a configuration, as done previously (14,36).

As an indicator for the enrichment of a given trait (e.g. belonging to the 3-state IPA model family or having a negatively regulated A→P transition) among a group of functional configurations (i.e. robust pulse filtering or pulse boosting configurations), an enrichment ratio was used. The enrichment ratio corresponds to the actual number of configurations with the trait among the functional configurations divided by the expected number of configurations with the trait among the same sized group under random selection. Specifically, the enrichment ratio for trait T, *E*_*T*_, was defined as:

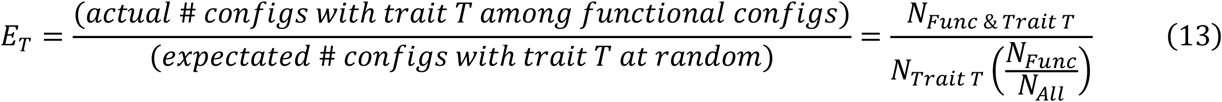

where *N*_*All*_ is the total number of configurations in the screen space, *N*_*Trait T*_ is the number of configurations in the screen space with trait T, *N*_*Func*_ is the number of functional configurations (i.e. robust pulse filtering or pulse boosting) in the screen space, and *N*_*Func*_ & _*Trait T*_ is the number of functional configurations with the trait T in the screen space. Note: If the trait T does not exist in the screen space (*N*_*Trait T*_ = 0) or there are no functional configurations in the screen space (*N*_*Func*_ = 0), then *E*_*T*_ is undefined. When investigating enrichment for transition types and transcription rate types, *E*_*T*_ was computed within the model family (2-state IA, 3-state IAR, 3-state IPA, or 4-state IPAR) because each model family has a different set of allowable transitions.

### Statistical Tests for Enrichment

To test the statistical significance of enrichment/depletion of a specific trait in a functional group (robust pulse filtering or boosting) with respect to its frequency in the entire screened set of configurations, the two-sided Fisher’s exact test was used (49). P values were corrected for multiple testing using the Benjamini-Hochberg procedure (50). In figures, [*] indicates an adjusted p-value < 0.05 and the lack of this symbol indicates a non-significant adjusted p-value.

### Gridded Parameter Sweeps and Variations in TF Dynamics

A subset of configurations containing features identified in Screen 1 were further subjected to gridded sweeps of all TF-unbound rate constants (*k*_*ij*,0_) from 0.01 to 100 min^-1^ on a logarithmic scale holding other parameters constant (Table S2: “Gridded Sweep, Base TF Dynamics”). Analyses conducted on these parameter sets were identical to those in Screens 1 and 2, except that *F*_*Continuous*_ ≥ 1.25 (instead of 2) was used for classification of activators to continuous inputs. In addition to base TF dynamics, these configurations were subjected to variations in TF dynamics at each parameter set above. The following variations of pulsatile TF dynamics parameterized by pulse on time (*τ*_*on*_), off time (*τ*_*off*_), and amplitude (*A*) were conducted: *τ*_*on*_ = (0.1, 0.316, 1, 3.16, 10 min) holding *τ*_*off*_ = 1 min and *A* = 10^8^ AU_1_ (saturating); *τ*_*off*_ = (0.1, 0.316, 1, 3.16, 10 min) holding *τ*_*on*_ = 1 min and *A* = 10^8^ AU_1_ (saturating); and *A* = (0.01, 0.1, 1, 10, 100 AU_1_) holding *τ*_*off*_ = 1 min and *τ*_*on*_ = 1 min (Table S2: “Gridded Sweep, Varied TF Dynamics”). Additionally, the following pulsatile TF dynamics parametrized by pulse frequency (*f*), duty cycle (*DC*), and amplitude (*A*) were conducted: *f* = (0.05, 0.158, 0.5, 1.58, 5 /min) holding *DC* = 0.5 and *A* = 10^8^ AU_1_ (saturating); *DC* = (0.10, 0.25, 0.50, 0.75, 0.90) holding *f* = 0.5/min and *A* = 10^8^ AU_1_ (saturating). Each was compared to sustained TF dynamics with a matched [TF] AUC, amplitude, and end time point.

### Computational Methods

Numerical solution of ODEs was performed in Python using the solve_ivp function with the LSODA method from the SciPy library. Latin hypercube samples were generated in Python using the SciPy library. Data visualization was conducted in Python using the Matplotlib library. Statistical tests were conducted in Python using the SciPy library.

## Data and Code Availability

All data reported in this paper will be shared by the lead contact, T. Curtis Shoyer, upon request. Code used to simulate and analyze mathematical models is available from the lead contact upon request and will be made publicly available at the time of publication.

## Acknowledgments

This project was supported by the Alexander von Humboldt-Stiftung (DE) through a Postdoctoral Fellowship awarded to TCS. We thank Dr. Edda Schulz and Dr. Trevor Ham for critical reading of the manuscript.

## Author contributions

TCS conceived the study, developed the model formulation and screen methodology, performed simulations, analyzed the data, wrote and edited the manuscript, and secured funding. BDV supervised the study, wrote and edited the manuscript, and secured funding.

## Declaration of generative AI and AI-assisted technologies in the manuscript preparation process

During the preparation of this work the authors used ChatGPT for language editing. After using this tool/service, the authors reviewed and edited the content as needed and take full responsibility for the content of the published article.

## Supplementary Information

Supplementary Information is provided as a separate document containing the following: Table S1-S2, Figures S1-S12 with legends, Supplementary Note 1, and References for Supplementary Information.

