## Supplementary Information for "Computational screen of promoter configurations that robustly sense transcription factor dynamics"

### Supplementary Tables

#### Table S1: Approximate Versions of Promoter State Models from Selected Experimental Literature on TF Dynamics Sensing and Refractoriness.

| Approximate Model and Classification (this paper) | | | | | Original Model and Experimental System (from citation) | | | |
|---|---|---|---|---|---|---|---|---|
| **ID** | **Approximate Model** | **Model Family** | **# Regulated Transitions** | **Simple / Bifunctional** | **Citation** | **TF** | **Target Gene(s)** | **Response to Sustained vs Pulsatile TF Dynamics** |
| 1 | 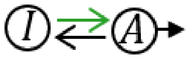 | 2-State IA | 1 | Simple | Chen et al. (1) | Crz1 | pYPS1-YFP and others | Pulse Boosting |
| 2 | 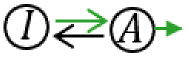 | 2-State IA | 1 | Simple | Hao and O'Shea (2) | Msn2 | Various | Pulse Filtering |
| 3 | 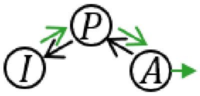 | 3-State IPA | 2 | Simple | Hansen and O'Shea (3) | Msn2 | Various | Pulse Filtering |
| 4 | 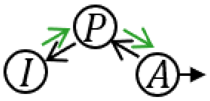 | 3-State IPA | 2 | Simple | Sweeney and McClean (4) | Msn2 | Various | Pulse Filtering |
| | | | | | Sen et al. (5) | NF-kB | Various | Not Compared in Study |
| 5 | 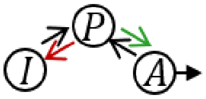 | 3-State IPA | 2 | Simple | Chen et al. (1) | Crz1 | pGYP7-YFP | Pulse Filtering |
| 6 | 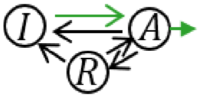 | 3-State IAR | 1 | Simple | Antwi et al. (6) | Synthetic TF | Synthetic Promoter/Reporter P4-iRFP670 and P5-iRFP670 | Pulse Filtering |
| 7 | 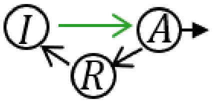 | 3-State IAR | 1 | Simple | Li et al. (7) | WCC | Candidate model for frq and vvd | Not Compared in Study |
| 8 | 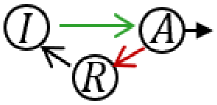 | 3-State IAR | 2 | Simple | Li et al. (7) | WCC | Candidate model for frq and vvd | Not Compared in Study |
| 9 | 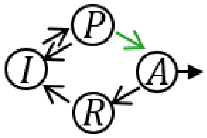 | 4-State IPAR | 1 | Simple | Li et al. (7) | WCC | Candidate model for frq and vvd | Not Compared in Study |

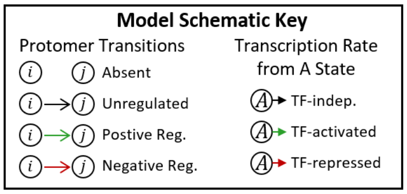

#### Table S2: Parameter Values.

| Symbol | Description | Screen 1 & 2 Value/Range | Screen 1 & 2 Distribution | Gridded Sweep, Base TF Dynamics | Gridded Sweep, Varied TF Dynamics |
|---|---|---|---|---|---|
| $\tau_{on}$ | TF Pulse On Time [min] | 1 | (Fixed value) | 1 | [0.1, 10] |
| $\tau_{off}$ | TF Pulse Off Time [min] | 1 | (Fixed value) | 1 | [0.1, 10] |
| $N_p$ | TF Pulse Number | 10 | (Fixed value) | 10 | 10 |
| $k_{ij,0}$ | TF-unbound Rate Constant for the $P_i$ to $P_j$ Promoter State Transition [1/min] | [0.001, 1000] | Log Uniform | [0.01, 100] | [0.01, 100] |
| $\epsilon_{ij}^{B}$ | Strength of TF Regulation for the $P_i$ to $P_j$ Promoter State Transition [no units] | [1,100] | Log Uniform | 10 | 10 |
| $k_{tr,max}$ | Maximum Regulated mRNA Transcription Rate [$AU_2$/min] | [0.1, 10] | Log Uniform | 1 | 1 |
| $\alpha_{tr,0}$ | Relative Basal Transcription ($k_{tr,basal} = \alpha_{tr,0} k_{tr,max}$) [no units] | 0.001 | (Fixed value) | 0.001 | 0.001 |
| $d_{mRNA}$ | mRNA Degradation Rate [1/min] | [0.1, 10] | Log Uniform | 0.1 | 0.1 |
| $A$ | TF Concentration [TF] during presence of TF [$AU_1$] | Non-saturating: 1<br>Saturating: $10^8$ | (Fixed value) | $10^8$ (Saturating) | Non-Saturating: [0.01, 100]<br>Saturating: $10^8$ |
| $K_{d,TF}$ | TF:RE Dissociation Constant [$AU_1$] | [0.01, 100] | Log Uniform | 1 | 1 |
| $n_{TF}$ | TF:RE Hill Coefficient [no units] | [0.5,4] | Uniform | 2 | 2 |

Note 1: $A$ and $K_{d,TF}$ are given in the same arbitrary units as [TF], which are defined as $AU_1$. [mRNA] is given in arbitrary units defined as $AU_2$. Note 2: Kinetic rates constants ($k_{ij,0}$, $k_{tr,max}$, and $d_{mRNA}$) were varied across ranges centered on the characteristic timescale of TF dynamics ($\tau_{on}$ and $\tau_{off}$). For simulations, $\tau_{on}$ and $\tau_{off}$ were set to the values given in this table (with units minutes).

#### Supplementary Figures

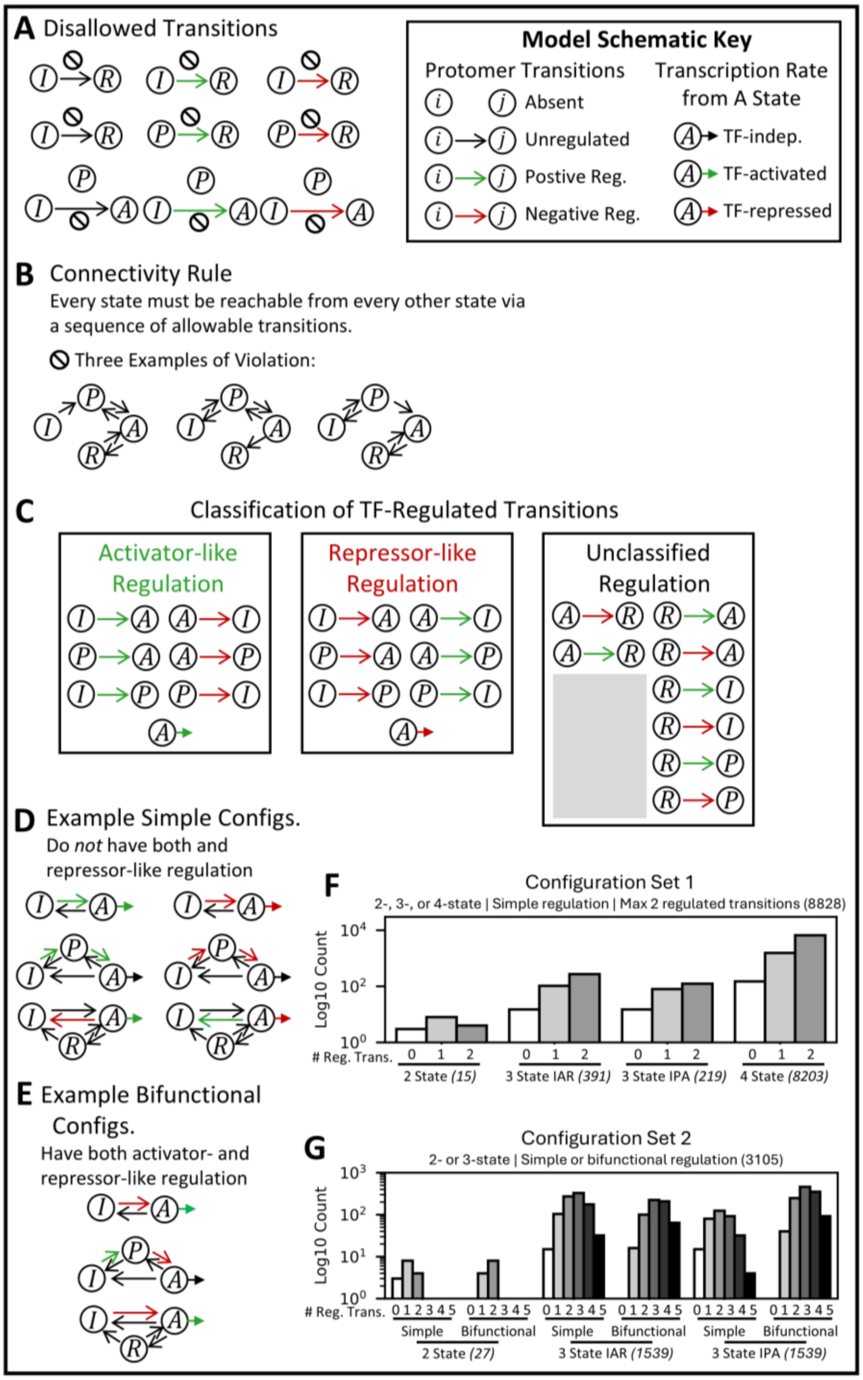


**Figure S1: Configuration enumeration and classification.** Related to Figure 1. A) Disallowed Transitions. B) Connectivity Rules. C) Classification of TF-regulated Transitions: Activator-like, Repressor-like, and Unclassified TF-based Regulation. D-E) Examples of Simple Configurations, which do not have both activator-like and repressor-like regulation, and Bifunctional Configurations, which have both activator-like and repressor-like regulation. F-G) Histograms of enumerated configurations indicating count (log scale) in each classification category for configuration set 1 (F) and configuration set 2 (G).

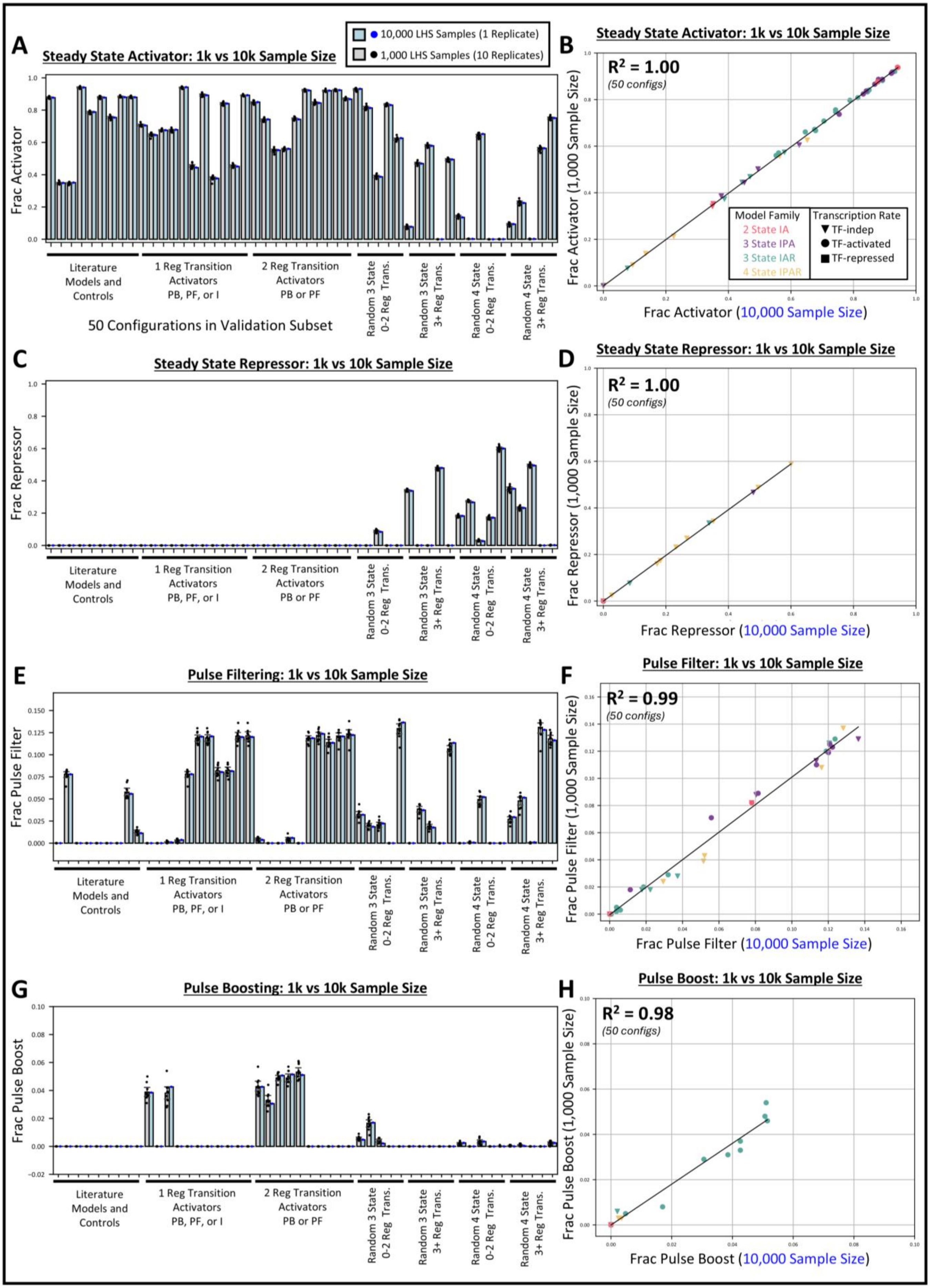

A
Steady State Activator: 1k vs 10k Sample Size
10,000 LHS Samples (1 Replicate)
1,000 LHS Samples (10 Replicates)
Frac Activator
Literature Models and Controls
1 Reg Transition Activators PB, PF, or I
2 Reg Transition Activators PB or PF
Random 3 State 0-2 Reg Trans.
Random 3 State 3+ Reg Trans.
Random 4 State 0-2 Reg Trans.
Random 4 State 3+ Reg Trans.
50 Configurations in Validation Subset
B
Steady State Activator: 1k vs 10k Sample Size
R² = 1.00
(50 configs)
Model Family
2 State IA
3 State IPA
3 State IAR
4 State IPAR
Transcription Rate
TF-indep
TF-activated
TF-repressed
Frac Activator (1,000 Sample Size)
Frac Activator (10,000 Sample Size)
C
Steady State Repressor: 1k vs 10k Sample Size
Frac Repressor
D
Steady State Repressor: 1k vs 10k Sample Size
R² = 1.00
(50 configs)
Frac Repressor (1,000 Sample Size)
Frac Repressor (10,000 Sample Size)
E
Pulse Filtering: 1k vs 10k Sample Size
Frac Pulse Filter
F
Pulse Filter: 1k vs 10k Sample Size
R² = 0.99
(50 configs)
Frac Pulse Filter (1,000 Sample Size)
Frac Pulse Filter (10,000 Sample Size)
G
Pulse Boosting: 1k vs 10k Sample Size
Frac Pulse Boost
H
Pulse Boost: 1k vs 10k Sample Size
R² = 0.98
(50 configs)
Frac Pulse Boost (1,000 Sample Size)
Frac Pulse Boost (10,000 Sample Size)

**Figure S2: Latin hypercube sampling with larger sample size and replicate samples.** Related to Figure 2. Screens using 1x independently drawn LHS sample of size 10,000 parameter sets and 10x independently drawn LHS samples of size 1,000 parameter sets were performed on a subset of 50 configurations chosen to represent variations in configuration properties (states and type and number of regulated transitions) and sensitivity to TF dynamics (PF: pulse filtering, PB: pulse boosting, and I: insensitive to dynamics). A) Fraction Activator Parameter Sets for 10x independently drawn LHS samples of size 1,000 (mean +/- 95% CI overlayed by a dot for each replicate) and 1x independently drawn LHS sample of size 10,000 for each of the 50 configurations. B) For the 50 configurations, scatter plot of Fraction Activator parameter sets for a LHS parameter set of size 1,000 (y axis) versus 10,000 (x axis). C-D) Same as (A-B), but for Fraction Repressor parameter sets. E-F) Same as (A-B), but for Fraction Pulse Filtering parameter sets (PSR<1/2). G-H) Same as (A-B), but for Fraction Pulse Boosting parameter sets (PSR>2).

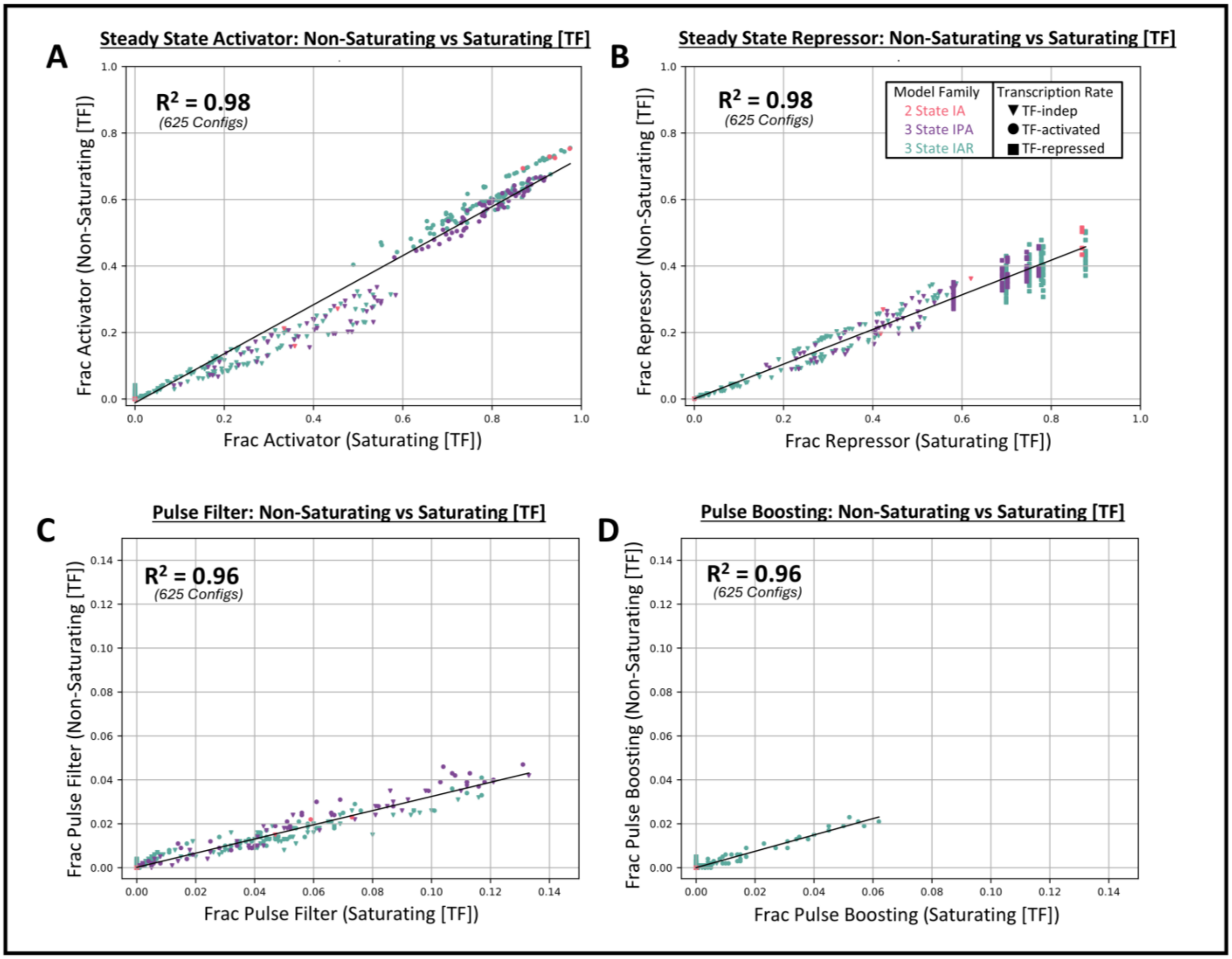


**Figure S3: Screening with saturating versus non-saturating TF concentration.** Related to Figure 2. For all 2- and 3-state configurations from Configuration set 1, comparison of (A) Fraction Activator parameter sets, (B) Fraction Repressor parameter sets, (C) Fraction Pulse Filtering parameter sets (PSR<1/2), and (D) Fraction Pulse Boosting parameter sets (PSR>2) for simulations with a saturating TF concentration ($A$=$10^8$; plotted on x-axis) versus simulations with a non-saturating TF concentration ($A$=1; plotted on y-axis).

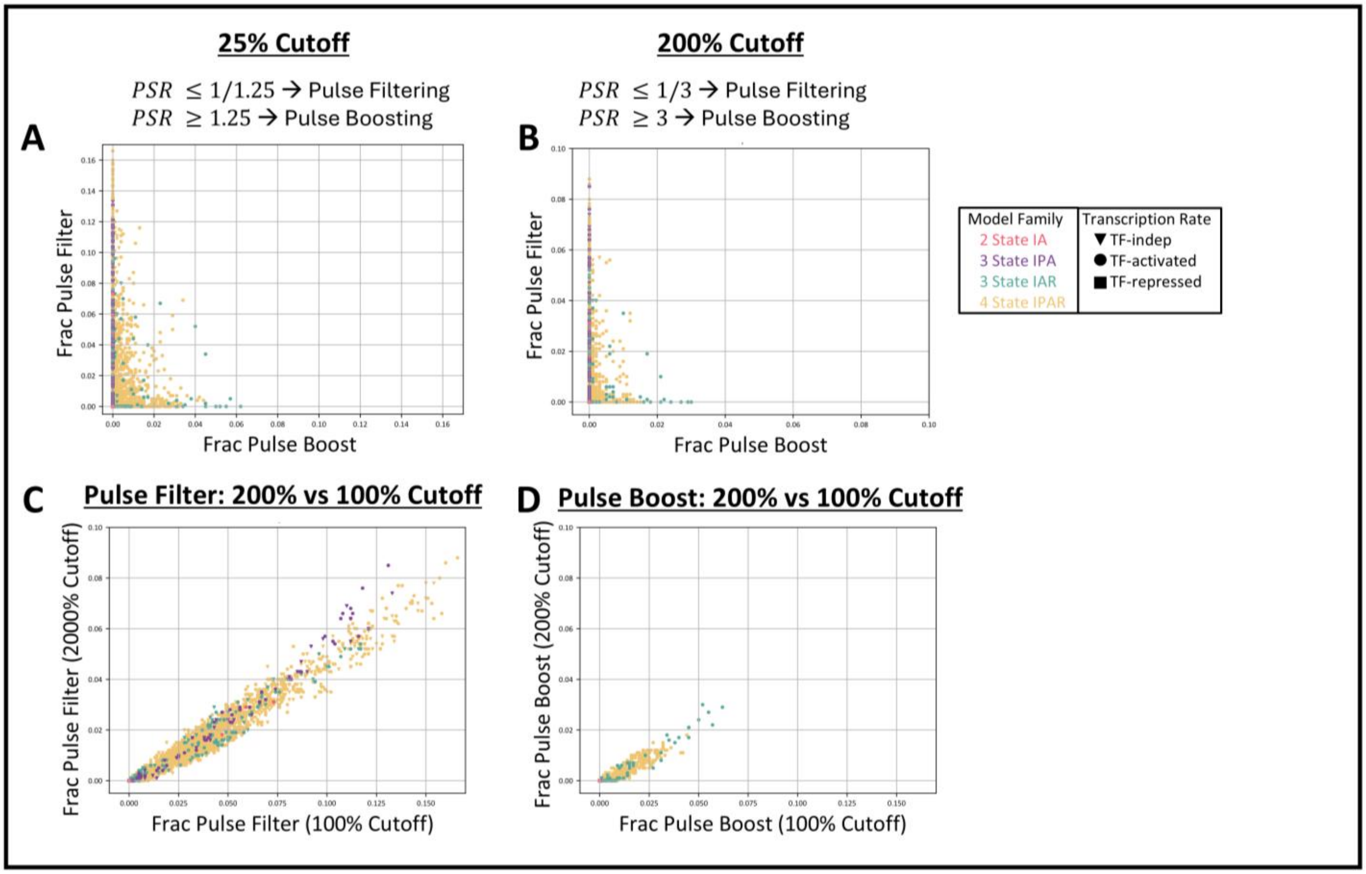


**Figure S4: Results from Screen 1 with lower or higher cutoffs for pulse filtering and boosting**. Related to Figure 2. A-B) Fraction Pulse Filtering parameter sets (Below Cutoff) vs Fraction Pulse Boosting parameter sets (Above Cutoff) for a lower (25%) and higher (200%) PSR Cutoff, compared to the normal PSR Cutoff of 100%. C) Scatter Plot of Fraction Pulse Filtering parameter sets using 200% Cutoff vs using 100% Cutoff. D) Same but for Fraction Pulse Boosting parameter sets. Note: The data are for the same configurations shown in Figure 2C.

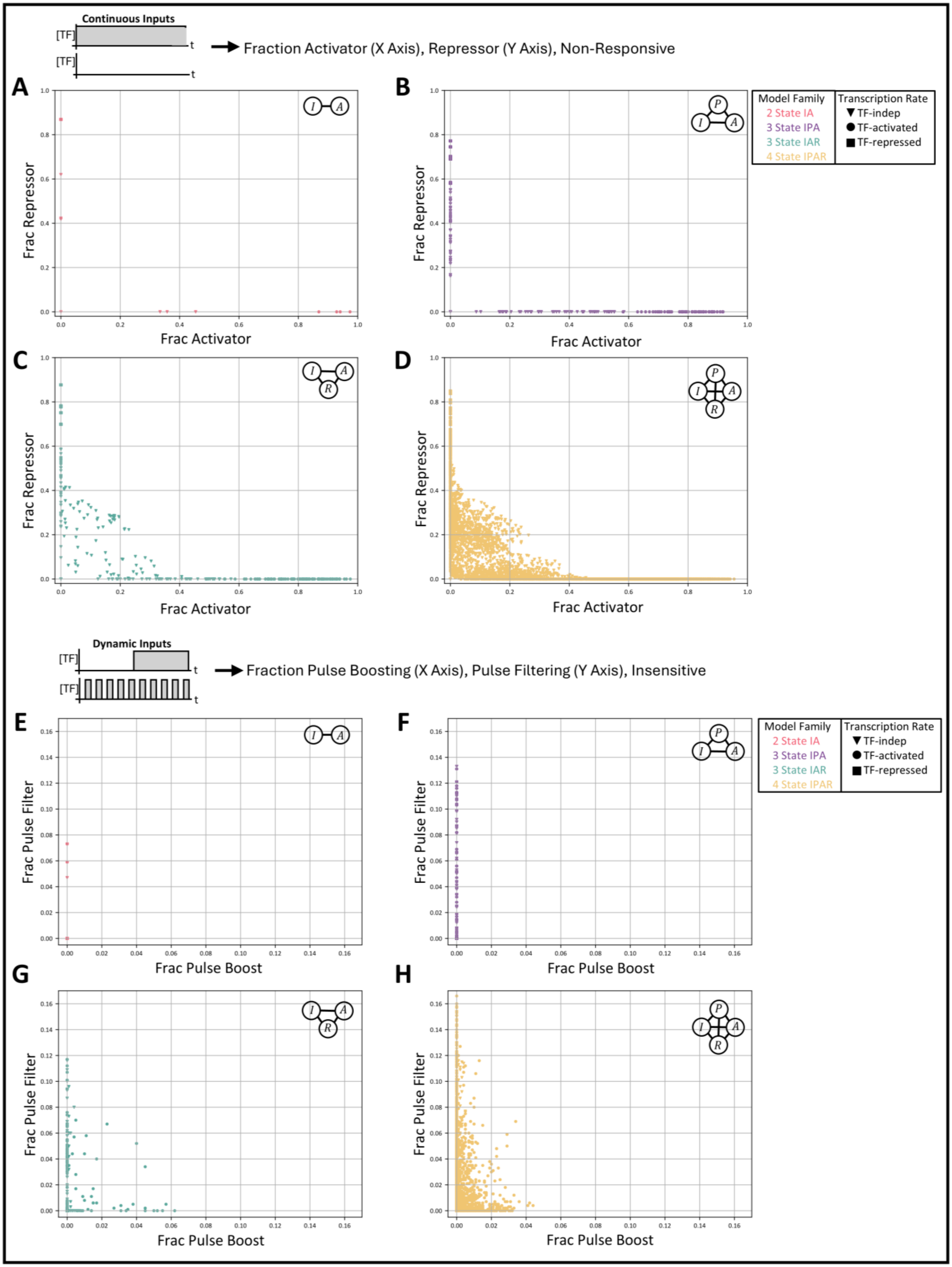

Continuous Inputs
[TF]
t
Fraction Activator (X Axis), Repressor (Y Axis), Non-Responsive
A
B
C
D
Frac Repressor
Frac Activator
Model Family
2 State IA
3 State IPA
3 State IAR
4 State IPAR
Transcription Rate
TF-indep
TF-activated
TF-repressed
Dynamic Inputs
Fraction Pulse Boosting (X Axis), Pulse Filtering (Y Axis), Insensitive
E
F
G
H
Frac Pulse Filter
Frac Pulse Boost

**Figure S5: Scatter plots for Screen 1 separated by model family.** Related to Figure 2. A-D) Response to continuous presence vs absence of TF shown as scatter plots of fraction of repressor parameter sets versus fraction of activator parameter sets, where each dot represents one configuration (1000 parameter sets). E-H) Response to sustained versus pulsatile TF dynamics shown as scatter plots of fraction pulse filtering parameter sets (PSR<1/2) versus fraction pulse boosting parameter sets (PSR>2), where each dot represents one topology (1000 parameter sets). Note: (E-H) contain the data from Figure 2C, separated by model family into different plots for better visualization.

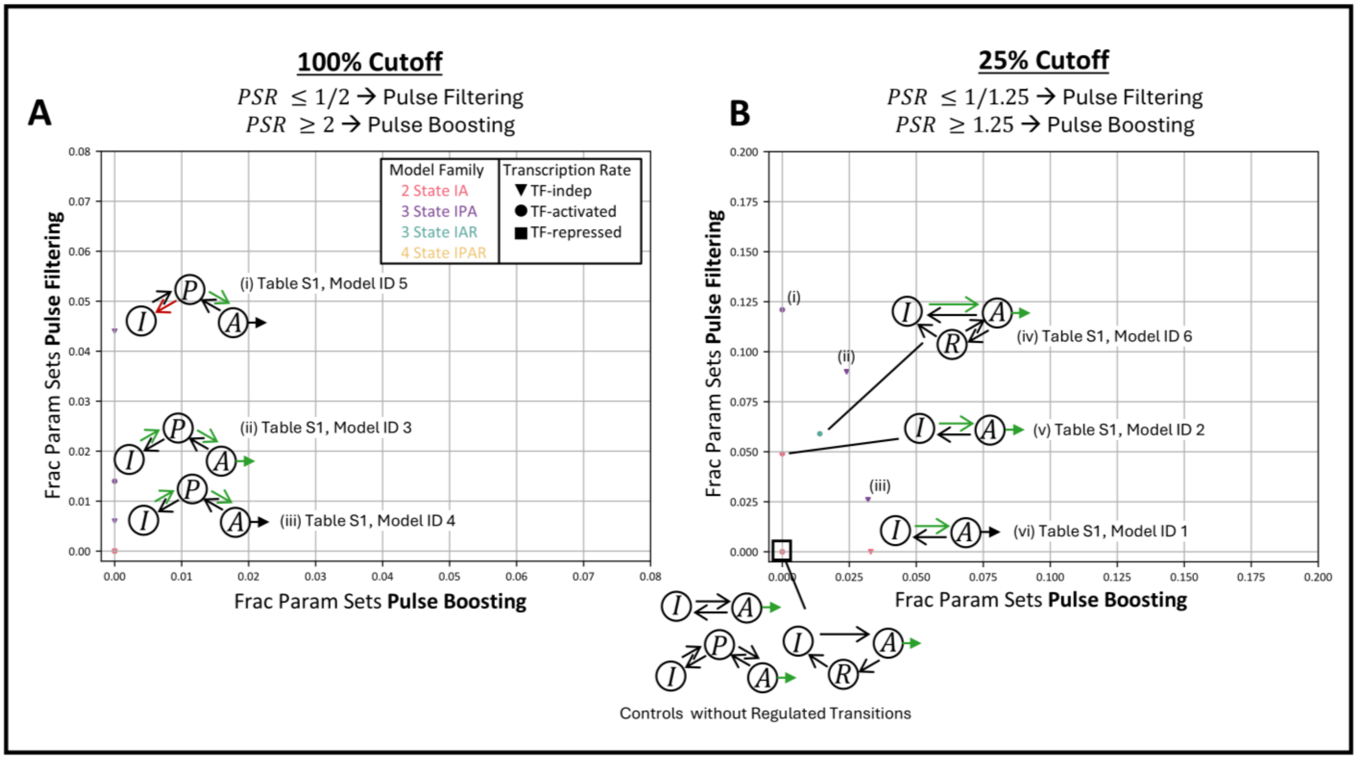


**Figure S6: Dynamics sensitivity of approximations to selected literature models contained in Screen 1.** Related to Figure 2. A) Scatter plot reproducing the data from Figure 2C for only a subset of configurations. Response to sustained versus pulsatile TF dynamics shown as scatter plots of fraction pulse filtering parameter sets (PSR<1/2) versus fraction pulse boosting parameter sets (PSR>2) sets, where each dot represents one topology (1000 parameter sets). B) Same as (A), but for a less stringent PSR cutoff indicated above the plot.

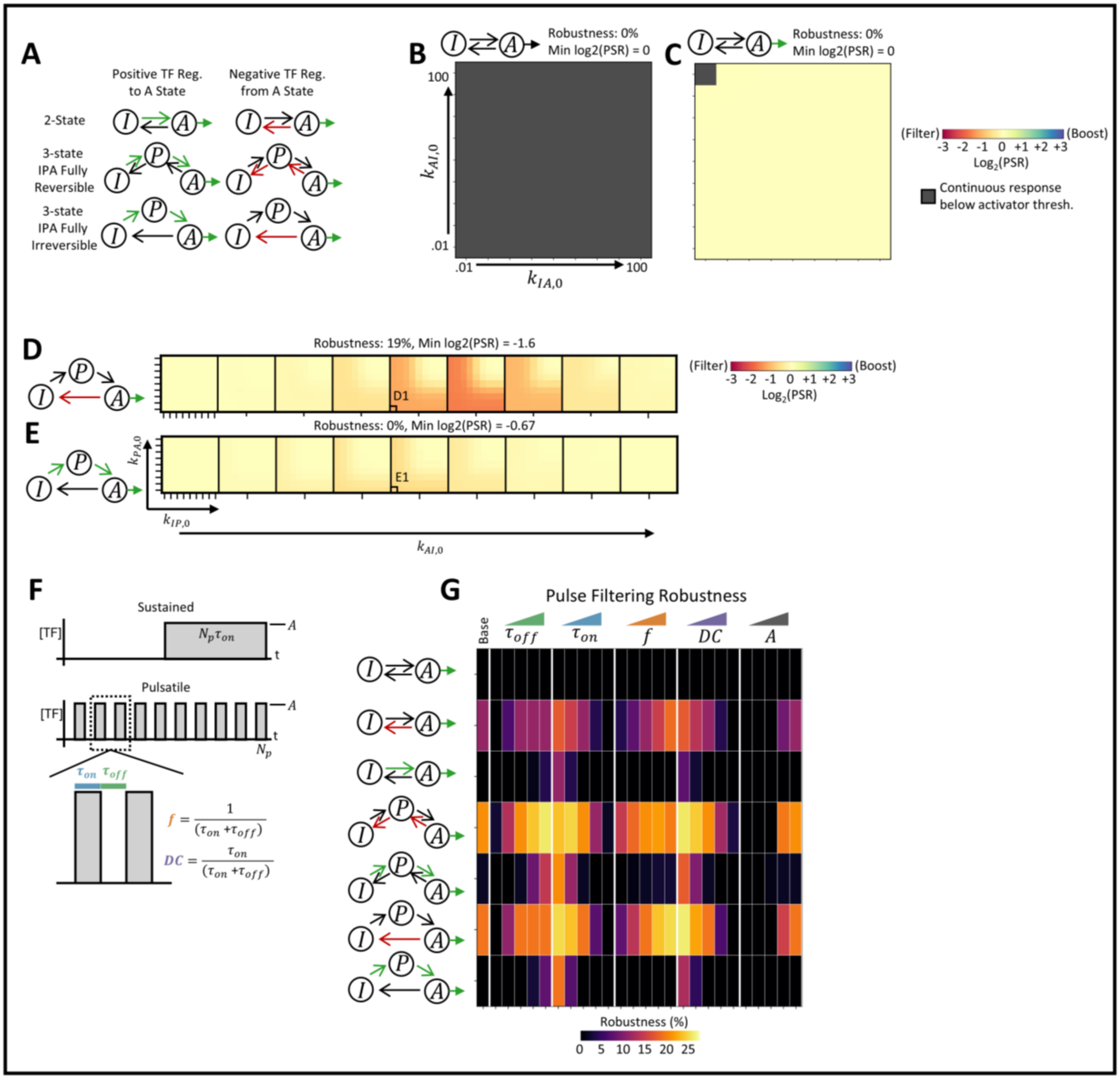


**Figure S7: Supporting analyses for pulse filtering gridded parameter sweeps and dynamics parameter sweeps**. Related to Figure 4. A) Schematics for a subset of 2-state and 3-state IPA models containing either positively regulated forward transitions (I→A or I→P→A) or negatively regulated reverse transitions (A→I or A→P→I). B-E) Heatmaps of log2 of PSR (negative/red indicates pulse filtering) for the indicated 2-state or 3-state IPA configurations showing gridded sweeps of the indicated TF-unbound rate constants (varied log uniformly from .01 to 100 $\mathrm{min}^{-1}$) with the base TF dynamics (Table S2: "Gridded Sweep, Base TF Dynamics"). Non-responsive parameter sets (less than 25% increase in mRNA in response to continuous presence vs absence of TF) are masked in grey. F) Schematic of pulsatile TF dynamics parameterized by pulse on time ($\tau_{on}$), off time ($\tau_{off}$), amplitude ($A$), or alternatively by pulse frequency ($f$) and duty cycle ($DC$). The corresponding sustained dynamics have a matched AUC, amplitude, and end time point. G) For the indicated configurations, pulse filtering robustness (fraction of parameter sets with PSR<1/2 and >25% increase in mRNA in response to continuous presence vs absence of TF) from gridded parameter sweeps performed at each of the following TF dynamics parameter sets: $\tau_{on}$ = (0.1, 0.316, 1,

3.16, 10 min) holding $\tau_{off}$ = 1 min and $A$ = $10^8$ $AU_1$ (saturating); $\tau_{off}$ = (0.1, 0.316, 1, 3.16, 10 min) holding $\tau_{on}$ = 1 min and $A$ = $10^8$ $AU_1$ (saturating); $f$ = (0.05, 0.158, 0.5, 1.58, 5 /min) holding $DC$ = 0.5 and $A$ = $10^8$ $AU_1$ (saturating); $DC$ = (0.10, 0.25, 0.50, 0.75, 0.90) holding $f$ = 0.5/min and $A$ = $10^8$ $AU_1$ (saturating); and $A$ = (0.01, 0.1, 1, 10, 100 $AU_1$) holding $\tau_{off}$ = 1 min and $\tau_{on}$ = 1 min (Table S2: "Gridded Sweep, Varied TF Dynamics").

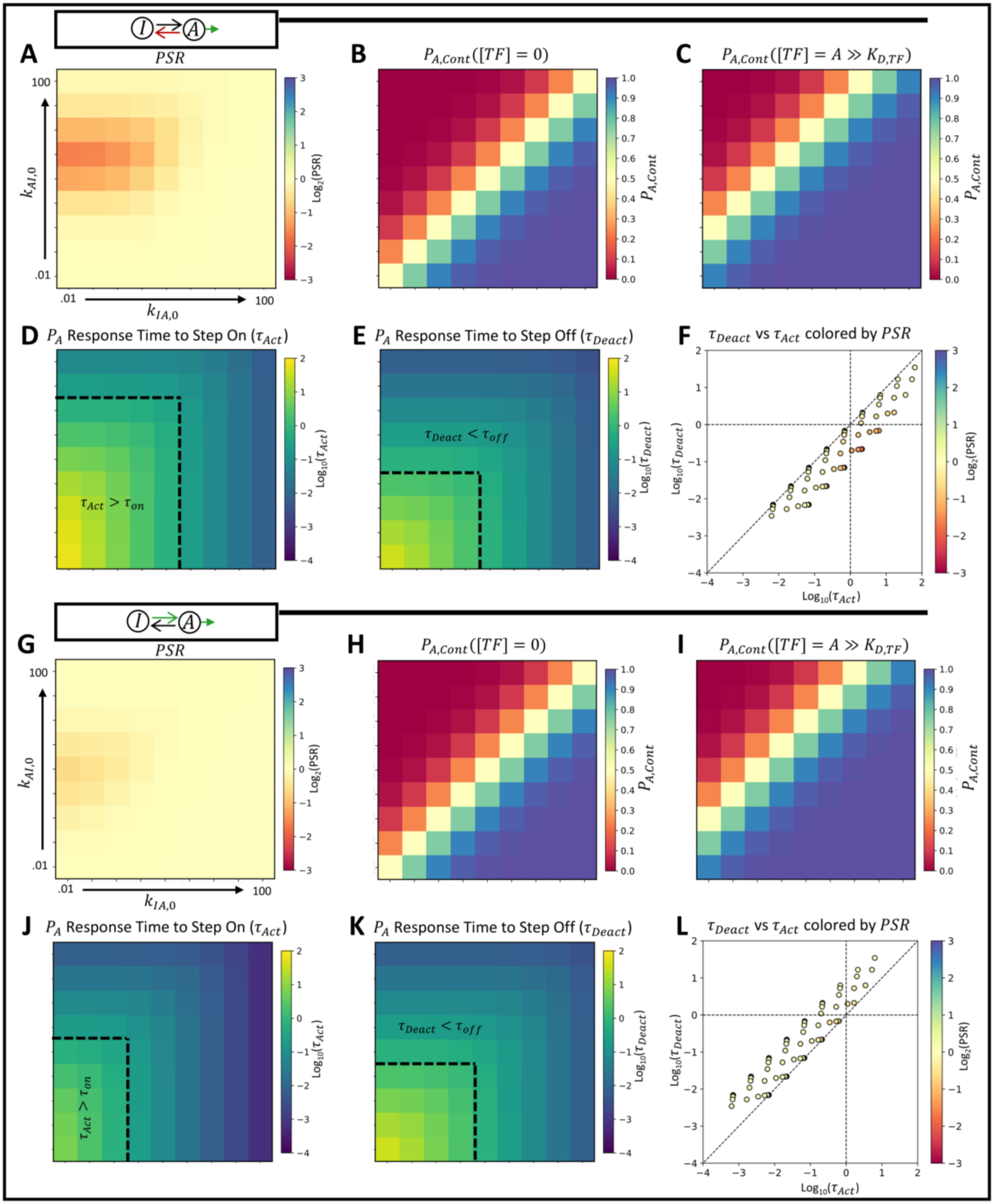


**Figure S8: Comparison of 2-state configurations with negatively regulated de-activation versus positively regulated activation.** Related to Figure 4. A) For the indicated 2-state configuration with negatively regulated A→I transition, heatmaps of log2 of PSR (negative/red indicates pulse filtering) showing gridded sweeps of the indicated TF-unbound rate constants (varied log uniformly from .01 to 100 min$^{-1}$) with the base TF dynamics (Table S2: "Gridded Sweeps, Base TF Dynamics"). B-E) Same as (A) but for the steady-state A state promoter in response to continuous absence of TF (B), steady-state A state promoter in response to continuous presence of TF (C), log10 of promoter activation half-time (time for half of the total increase in A state in response to a step change from absence to presence of TF) (D), log10 of the promoter deactivation half-time (time for half of the total decrease in A state in response to a step change from presence to absence of TF) (E). F) Scatter plot of log10 promoter deactivation half-time vs log10 promoter activation half-time colored by log2(PSR). G-L) For the indicated 2-state configuration with positively regulated I→A transition, same as (A-F).

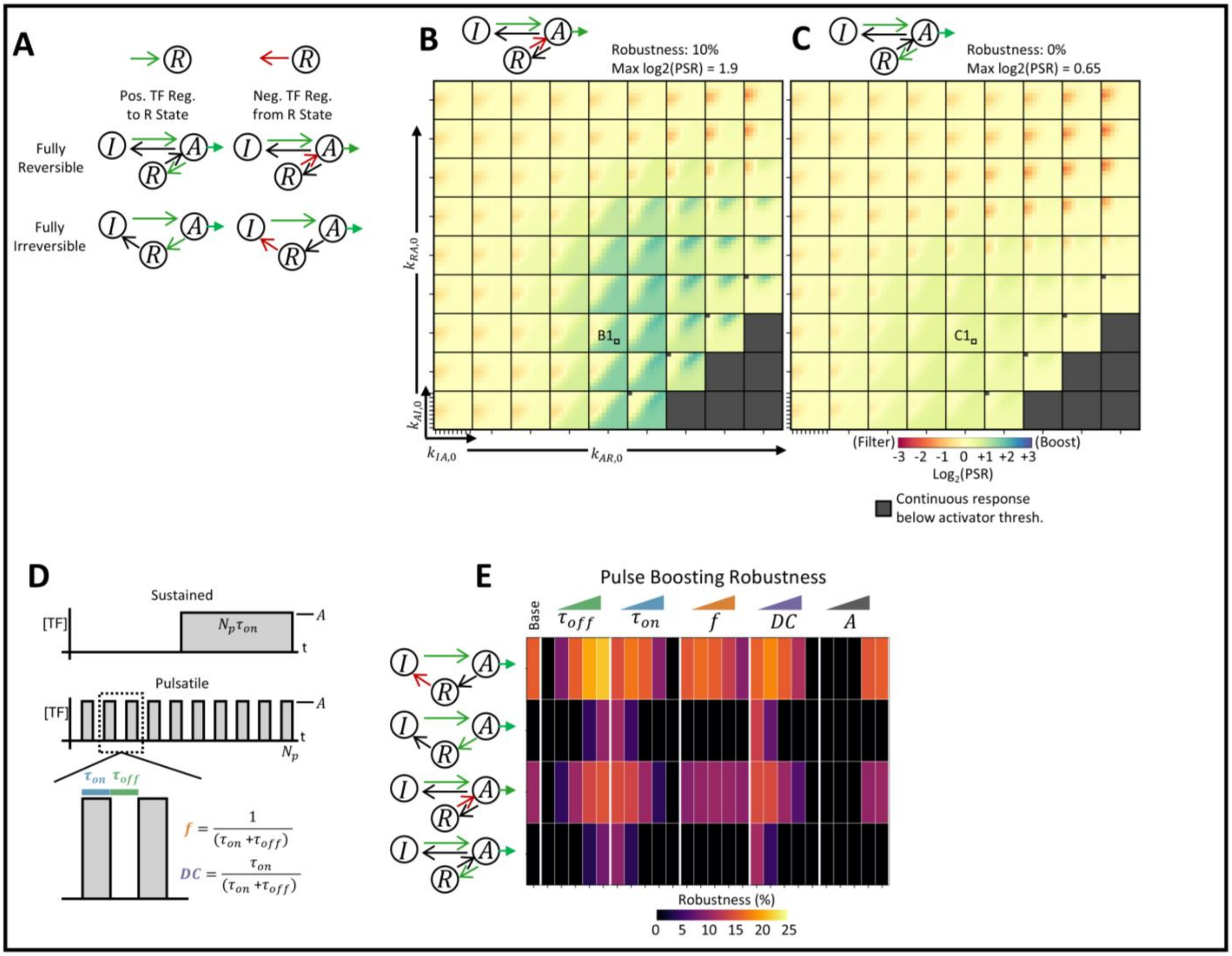


**Figure S9: Supporting analyses for pulse boosting gridded parameter sweeps and dynamics parameter sweeps**. Related to Figure 6. A) Schematics of a subset of configurations containing fully irreversible and fully reversible versions of the 3-state IAR model with either positive regulation of transitions into the R state or negative regulation of transitions out of the R state. B-C) Heatmaps of log2 of PSR (positive/blue indicates pulse boosting) for the indicated 2-state or 3-state IPA configurations showing gridded sweeps of the indicated TF-unbound rate constants (varied log uniformly from .01 to 100 $min^{-1}$) with the base TF dynamics (Table S2: “Gridded Sweep, Base TF Dynamics”). Non-responsive parameter sets (less than 25% increase in mRNA in response to continuous presence vs absence of TF) are masked in grey. D) Schematic of pulsatile TF dynamics parameterized by pulse on time ($\tau_{on}$), off time ($\tau_{off}$), amplitude ($A$), or alternatively by pulse frequency ($f$) and duty cycle ($DC$). The corresponding sustained dynamics have a matched AUC, amplitude, and end time point. E) For the indicated configurations, pulse boosting robustness (fraction of parameter sets with PSR>2 and >25% increase in mRNA in response to continuous presence vs absence of TF) from gridded parameter sweeps performed at each of the following TF dynamics parameter sets: $\tau_{on}$ = (0.1, 0.316, 1, 3.16, 10 min) holding $\tau_{off}$ = 1 min and $A$ = $10^8$ $AU_1$ (saturating); $\tau_{off}$ = (0.1, 0.316, 1, 3.16, 10 min) holding $\tau_{on}$ = 1 min and $A$ = $10^8$ $AU_1$ (saturating); $f$ = (0.05, 0.158, 0.5, 1.58, 5 /min) holding $DC$ = 0.5 and $A$ = $10^8$ $AU_1$ (saturating); $DC$ = (0.10, 0.25, 0.50, 0.75, 0.90) holding $f$ = 0.5/min and $A$ = $10^8$ $AU_1$ (saturating); and $A$ = (0.01, 0.1, 1, 10, 100 $AU_1$) holding $\tau_{off}$ = 1 min and $\tau_{on}$ = 1 min (Table S2: “Gridded Sweep, Varied TF Dynamics”).

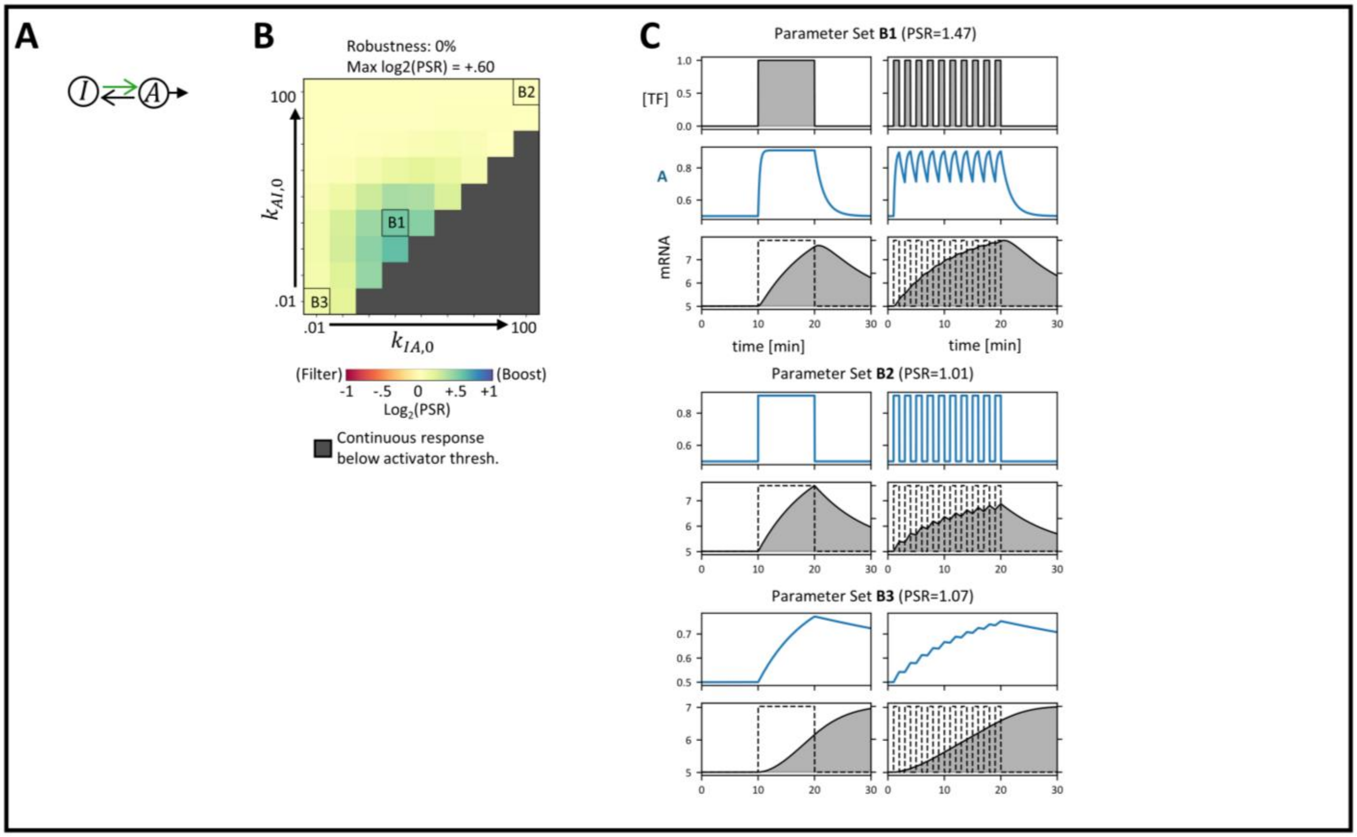


**Figure S10: Weak pulse boosting in a 2-state model.** Related to Figure 6. A) Schematic of promoter configuration. B) Heatmap of log2 of PSR (positive/blue indicates pulse boosting) for the indicated configuration showing gridded parameter sweeps of the indicated TF-unbound rate constants (varied log uniformly from .01 to 100 $\mathrm{min}^{-1}$) with the base TF dynamics (Table S2: "Gridded Sweep, Base TF Dynamics"). Parameter sets with less than 25% increase in mRNA in response to continuous presence vs absence of TF are considered Non-Responsive and masked in grey. C) For the indicated parameter sets from (B), time courses of the following for sustained (left) and pulsatile (right) inputs on matched axes: [TF] ($\times 10^8$ $AU_1$; shown for first parameter set only), A state fraction, and mRNA level ($AU_2$) versus time (min). Only shown up to 10 min post-stimulation.

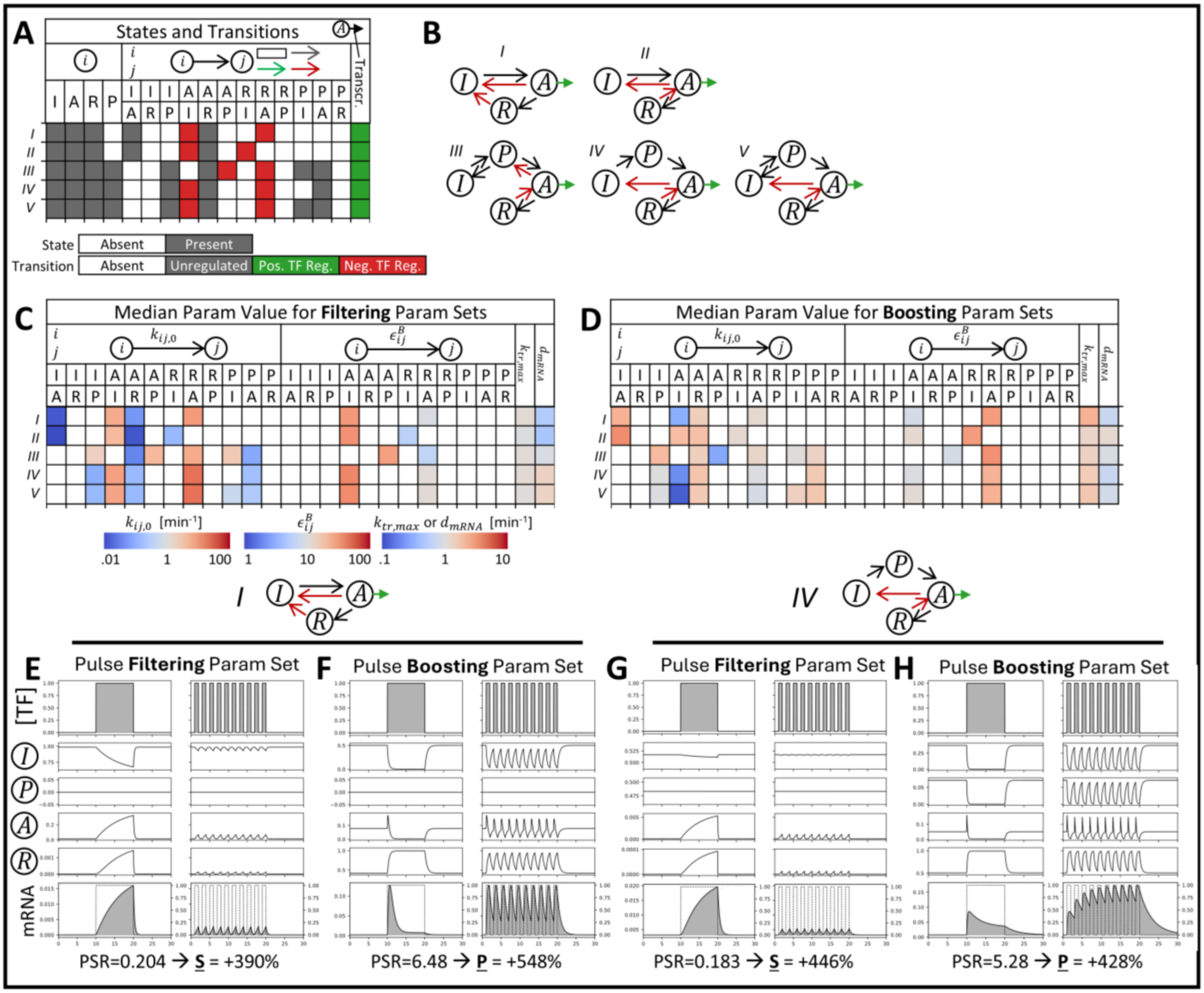

**Figure S11: Configurations that perform both pulse filtering and pulse boosting.** Related to Figure 2. A-B) All configurations having at least 2.5% parameter sets classified as pulse filtering (PSR<1/2) and at least 2.5% parameter sets classified as pulse boosting (PSR>2). Each row in the table corresponds to a configuration with the columns indicating the identity of the configuration: states, transitions, transcription rate type. C) Median parameter values across all pulse filtering parameter sets (unvaried parameters shown in white). Each row is a configuration from (A-B). D) Same for pulse boosting parameter sets. E-F) For configuration I, time courses for pulse filtering and pulse boosting parameter sets, respectively, each showing the following for sustained (left) and pulsatile (right) inputs on matched axes: [TF] (x$10^8$ $AU_1$), promoter state fractions (I/P/A/R), and mRNA level ($AU_2$) versus time (min). PSR given below. Only shown up to 10 min post-stimulation. G-H) Same for configuration IV. Note: The configurations in this plot come from the scatter plots in Figure 2C.

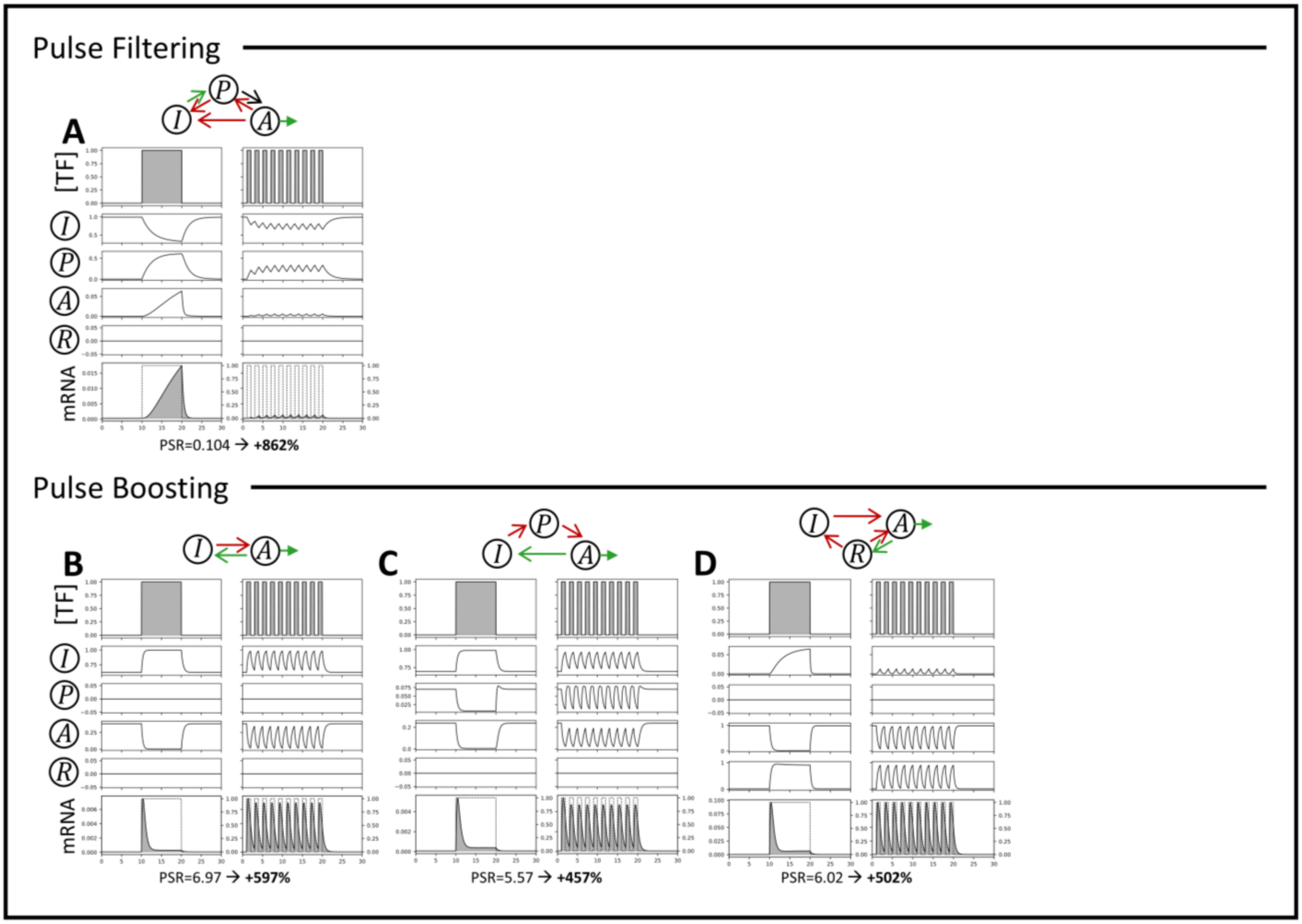


**Figure S12: Representative time courses for pulse filtering and boosting configurations in Screen 2**. Related to Figure 7. Time courses for representative configuration/parameter sets performing (A) pulse filtering or (B-D) pulse boosting, showing the following for sustained (left) and pulsatile (right) inputs on matched axes: [TF] (x$10^8$ $AU_1$), promoter state fractions (I/P/A/R), and mRNA level ($AU_2$) versus time (min). Only shown up to 10 min post-stimulation. PSR given below.

### Supplementary Note 1

#### Parameters for time courses in Figure 3:

(B) $k_{\mathrm{IP},0}$: 0.51; $k_{\mathrm{PI},0}$: 85; $k_{\mathrm{PA},0}$: .0074; $k_{\mathrm{AP},0}$: 1.3; $1/\epsilon_{AP}^{B}$: .013; $1/\epsilon_{PI}^{B}$: .030; $k_{\mathrm{tr},max}$: .49; $d_{\mathrm{mRNA}}$ : 9.3

(C) $k_{\mathrm{IP},0}$: .0064; $k_{\mathrm{PI},0}$: .82; $k_{\mathrm{PA},0}$: .0011; $k_{\mathrm{AP},0}$: 9.6; $1/\epsilon_{AP}^{B}$: .010; $1/\epsilon_{PI}^{B}$: .054; $k_{\mathrm{tr},max}$: .26; $d_{\mathrm{mRNA}}$ : 3.7

(D) $k_{\mathrm{IP},0}$: .060; $k_{\mathrm{PI},0}$: 11; $k_{\mathrm{PA},0}$: .048; $k_{\mathrm{AR},0}$: 1.9; $k_{\mathrm{RI},0}$: .20; $1/\epsilon_{AR}^{B}$: .44; $1/\epsilon_{PI}^{B}$: .019; $k_{\mathrm{tr},max}$: .65; $d_{\mathrm{mRNA}}$ : 2.7

(E) $k_{\mathrm{IP},0}$: 1.3; $k_{\mathrm{PI},0}$: 14; $k_{\mathrm{PA},0}$: .024; $k_{\mathrm{AR},0}$: 1.7; $k_{\mathrm{RP},0}$: 160; $1/\epsilon_{AR}^{B}$: .028; $1/\epsilon_{PI}^{B}$: .23; $k_{\mathrm{tr},max}$: .16; $d_{\mathrm{mRNA}}$ : .38

Units for rate constants are min$^{-1}$ and $\epsilon_{ij}^{B}$ parameters are unitless. For positive regulation, $\epsilon_{ij}^{B}$ is given. For negative regulation, $1/\epsilon_{ij}^{B}$ is given. $K_{d,TF}$ and $n_{TF}$ are not reported because the saturating TF regime is used. Configurations are given in Figure 3 by the schematics above each time course.

#### Parameters for time courses in Figure 5:

(B) $k_{IA.0}$: .65; $k_{\mathrm{AR},0}$: 5.9; $k_{\mathrm{RI},0}$: 11; $\epsilon_{IA}^{B}$: 37; $1/\epsilon_{RI}^{B}$: .014; $k_{\mathrm{tr},max}$: .30; $d_{\mathrm{mRNA}}$ : 7.2

(C) $k_{IA.0}$: .37; $k_{\mathrm{AR},0}$: 1.1; $k_{\mathrm{RI},0}$: .15; $\epsilon_{IA}^{B}$: 24; $1/\epsilon_{RI}^{B}$: .023; $k_{\mathrm{tr},max}$: 1.7; $d_{\mathrm{mRNA}}$ : 7.4

(D) $k_{IA.0}$: .98; $k_{\mathrm{AR},0}$: 1.6; $k_{\mathrm{RI},0}$: .16; $\epsilon_{IA}^{B}$: 2.9; $1/\epsilon_{RI}^{B}$: .012; $k_{\mathrm{tr},max}$: .40; $d_{\mathrm{mRNA}}$ : .20

(E) $k_{IP.0}$: 510; $k_{\mathrm{PA},0}$: .73; $k_{\mathrm{AI},0}$: 29; $k_{\mathrm{AR},0}$: 5.7; $k_{\mathrm{RA},0}$: 1.1; $\epsilon_{PA}^{B}$: 6.1; $1/\epsilon_{RI}^{B}$: .030; $k_{\mathrm{tr},max}$: 2.4; $d_{\mathrm{mRNA}}$ : 1.1

Units for rate constants are min$^{-1}$ and $\epsilon_{ij}^{B}$ parameters are unitless. For positive regulation, $\epsilon_{ij}^{B}$ is given. For negative regulation, $1/\epsilon_{ij}^{B}$ is given. $K_{d,TF}$ and $n_{TF}$ are not reported because the saturating TF regime is used. Configurations are given in Figure 5 by the schematics above each time course.
